# Mechanisms of KCNQ1 gating modulation by KCNE1/3 for cell-specific function

**DOI:** 10.1101/2025.06.08.658478

**Authors:** Chenxi Cui, Lu Zhao, Ali A. Kermani, Shuzong Du, Tanadet Pipatpolkai, Meiqin Jiang, Sagar Chittori, Yong Zi Tan, Jingyi Shi, Lucie Delemotte, Jianmin Cui, Ji Sun

## Abstract

KCNQ1 potassium channels are essential for physiological processes such as cardiac rhythm and intestinal chloride secretion. KCNE-family subunits (KCNE1-5) associate with KCNQ1, conferring distinct properties across various tissues. KCNQ1 activation requires membrane depolarization and phosphatidylinositol 4,5-bisphosphate (PIP2) whose cellular levels are controlled by Gαq-coupled GPCR activation. While modulation of KCNQ1’s voltage-dependent activation by KCNE1/3 is well-characterized, their effects on PIP2-dependent gating of KCNQ1 *via* GPCR signaling remain less understood. Here we resolved structures of KCNQ1–KCNE1 and reassessed reported KCNQ1-KCNE3 structures with and without PIP2. We revealed that KCNQ1–KCNE1/3 complexes feature two PIP2-binding sites, with KCNE1/3 contributing to a previously overlooked, uncharacterized site involving residues critical for voltage sensor and pore domain coupling. Via this site, KCNE1 and KCNE3 distinctly modulate the PIP2-dependent gating, in addition to the voltage sensitivity, of KCNQ1. Consequently, KCNE3 converts KCNQ1 into a voltage-insensitive PIP2-gated channel governed by GPCR signaling to maintain ion homeostasis in non-excitable cells. KCNE1, by significantly enhancing KCNQ1’s PIP2 affinity and resistance to GPCR regulation, forms predominantly voltage-gated channels with KCNQ1 for conducting the slow-delayed rectifier current in excitable cardiac cells. Our study highlights how KCNE1/3 modulates KCNQ1 gating in different cellular contexts, providing insights for tissue-specifically targeting multi-functional channels.

## INTRODUCTION

KCNQ1, also known as KvLQT1 or Kv7.1, is a voltage- and lipid-gated potassium channel, found in various organs such as the heart, inner ear, colon, kidney, pancreas and stomach^1-8^. It plays a vital role in physiological processes including cardiac rhythm regulation, auditory function, and gastric acid secretion^9^. Mutations in KCNQ1 protein or pharmacological interference with its channel function can result in disorders like long-QT syndromes, short-QT syndromes, atrial fibrillation, diabetes mellitus, and deafness, highlighting its potential as a drug target^10-12^.

KCNQ1 forms channel complexes with KCNEs (KCNE1–5), a family of single transmembrane proteins that function as beta subunits for KCNQ1, modifying its voltage-dependent gating behavior and conferring cell-specific biophysical properties^1,13-15^. In excitable cells such as cardiomyocytes, KCNQ1 associates with KCNE1, which alters its voltage-dependent activation by slowing activation and deactivation kinetics, shifting voltage dependence, increasing single-channel conductance, and preventing inactivation (Figure 1A). These properties enable the KCNQ1–KCNE1 complex to conduct the vital slow-delayed rectifier current (*I_Ks_*) during the repolarization phase of cardiac action potentials^7,8,16,17^. In contrast, in non-excitable cells such as polarized epithelial cells in the colon, small intestine, and airways^6,18-20^, KCNE3 renders KCNQ1 voltage-insensitive, with the channel complex exhibiting a linear current-voltage relationship at physiological membrane potentials (Figure 1A). How this voltage-insensitive KCNQ1–KCNE3 channel complex is gated remains to be clarified, and it has been thought as being constitutively open, a behavior that would facilitate potassium flow to support epithelial chloride secretion^2,6,21^ but could also be disruptive if uncontrolled. The other KCNE subunits (KCNE2, KCNE4, and KCNE5) can regulate and alter the channel properties of KCNQ1^22-24^, but their precise physiological functions are less understood.

**Figure 1:**
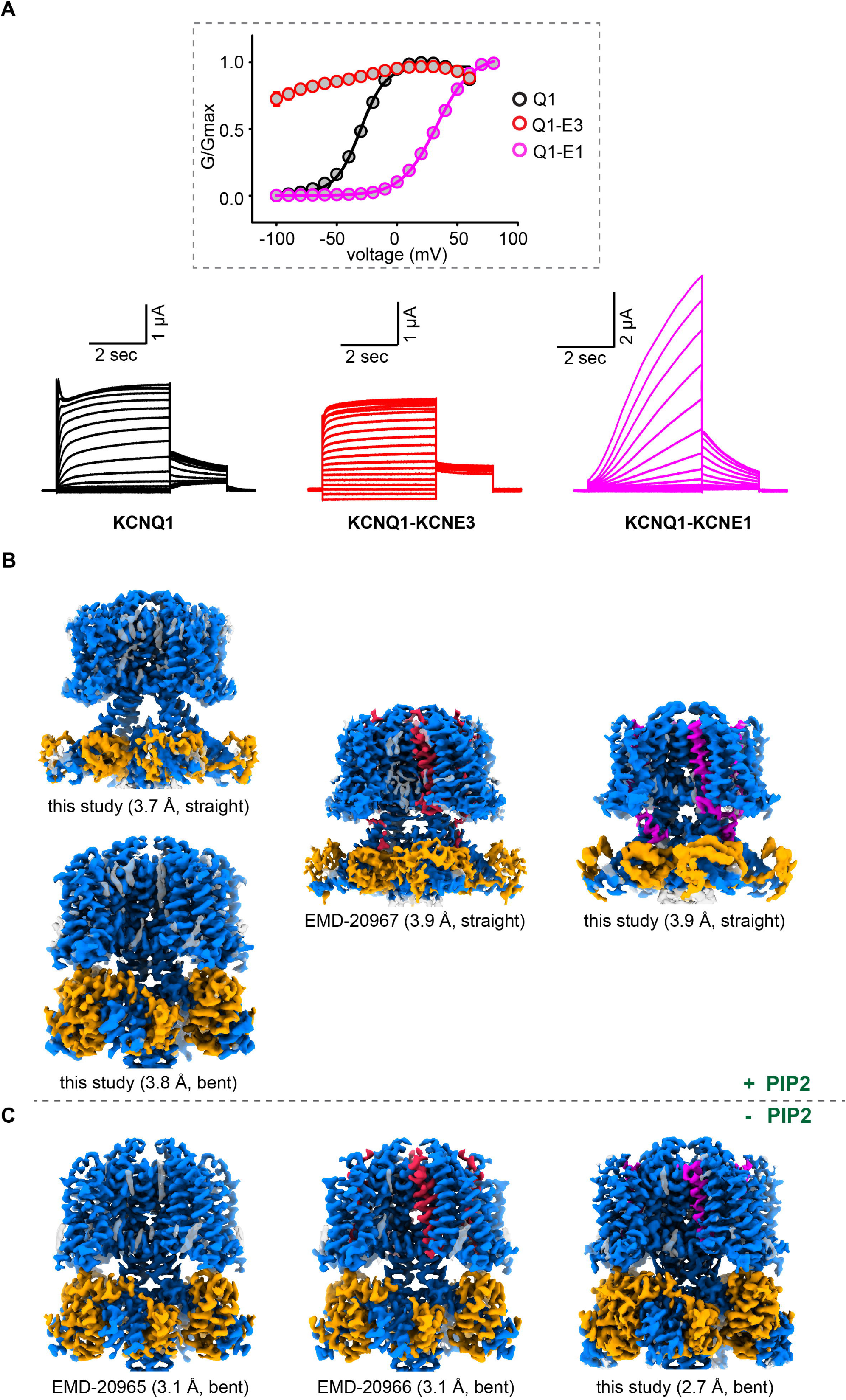
Structural and functional characterization of KCNQ1 and KCNQ1-KCNEs. **A.** Electrophysiological recording curves of KCNQ1 (black), KCNQ1–KCNE3 (red) and KCNQ1– KCNE1 (magenta) in oocytes. The G-V curve (upper panel) is plotted in the inset using the same color code as the raw traces (lower panel). In lower panel, the left (black), middle (red) and right (magenta) are representative electrophysiological traces of KCNQ1 wild type (WT), KCNQ1–KCNE3, KCNQ1-KCNE1 complexes, respectively. Voltage steps are applied from −100 mV to 60 mV for KCNQ1 and KCNQ1–KCNE3 at 10-mV increments, and −100 mV to 80 mV for KCNQ1–KCNE1 at 10-mV increments. Holding voltage is -80 mV for 0.5 s and tail current was recorded at -40 mV for 2 s (n ≥10), All constructs used are wild-type full-length proteins in the electrophysiology recording. **B.** Cryo-EM maps of KCNQ1 (left), KCNQ1–KCNE3 (middle) and KCNQ1–KCNE1 (right) in nanodiscs with the presence of PIP2. **C.** Cryo-EM maps of KCNQ1, KCNQ1–KCNE3 and KCNQ1-KCNE1 in nanodiscs with the absence of PIP2. In **B–C**, KCNQ1, CaM, KCNE1 and KCNE3 are colored in blue, orange, magenta, and red respectively. Overall resolutions of the cryo-EM maps are indicated. Constructs used for cryo-EM analysis are described in the method session with N- and C-terminus loop truncation.

The potassium conductance of KCNQ1 or KCNQ1–KCNE1/3 channel complexes require PIP2^25-30^. PIP2 is a key signaling lipid in the Gαq protein-coupled receptor (GqPCR) signaling pathway, where GPCR activation leads to PIP2 hydrolysis by phospholipase C (PLC)^31^. Notably, KCNQ2–KCNQ5 channels, neuronal paralogs of KCNQ1, conduct the classical M currents^32^, for they are controlled by Gαq-coupled muscarinic acetylcholine receptors. Yet, the modulation of KCNQ1 and its KCNE complexes by GPCR pathways remains less understood. We previously showed that KCNE1 enhances KCNQ1’s affinity for PIP2^33^, but it is unclear whether KCNE3 has a similar effect and whether (and how) KCNQ1–KCNE1/3 responds to GPCR stimulations. More importantly, how homologous KCNE1 and KCNE3 tune the voltage- and lipid-dependent gating properties of KCNQ1 to accommodate its cell-specific physiological roles remains elusive.

Here we explore these open questions to understand how KCNE1 and KCNE3 modulate the voltage and lipid dependencies of KCNQ1 to support its functions in excitable and non-excitable cells. We present high-resolution structures of KCNQ1 (+PIP2) and KCNQ1–KCNE1 (+/-PIP2) (Figure 1B-C). We integrated those data and previous structures of KCNQ1 (-PIP2)^28^ and KCNQ1–KCNE3 (+/-PIP2)^29^ with electrophysiology assays and molecular dynamics (MD) simulations. Notably, we revealed that structures of active KCNQ1 and its KCNE1/3 complexes feature a second PIP2-binding site, in addition to the one we previously reported^29^. Like KCNE1^33^, KCNE3, modifies the PIP2-binding affinity of KCNQ1, but to a different extent. The lower affinity renders KCNQ1–KCNE3 more sensitive to PLC activation and GqPCR regulation than KCNQ1–KCNE1. Therefore, the KCNQ1–KCNE3 complex is a PIP2-gated channel, regulated by GPCR signaling. Taken together, we reveal an unexpected channel regulation mechanism where KCNQ1 functions as a predominantly voltage-gated or a PIP2-gated channel by association with either KCNE1 or KCNE3 subunits, thus adapting to different cellular contexts.

## RESULTS

### Structural characterization of KCNQ1 and KCNQ1–KCNE1/3 complexes

We set out to investigate structures of the KCNQ1–KCNE1 complex. Unlike KCNQ1–KCNE3^29^, KCNQ1–KCNE1 does not form a stable complex in detergent solutions. To address this issue, we introduced a cysteine mutation pair to their extracellular regions, as previously described^34^. The resulting complex (KCNQ1^I145C^–KCNE1^K41C^) showed biochemical stability and biophysical properties nearly identical to the wild-type complex (Figures S1A–S1D), enabling structural determination of this weakly associated complex (Figures S1E–S1G). The KCNQ1^I145C^– KCNE1^K41C^ complex was stable during purification in the absence of reducing reagents, but they did not appear to form a disulfide bond in the cryo-electron microscope (cryo-EM) structure (Figure S1H).

Same as KCNQ1–KCNE3 (+/-PIP2) complexes^29^, we prepared samples in lipid nanodiscs^29,35,36^ for cryo-EM characterization of human KCNQ1 (+PIP2) and KCNQ1–KCNE1 (+/-PIP2) channel complexes and obtained near-atomic resolution structures. The samples were frozen under depolarization conditions, so voltage sensors are in the activated “up” state (Figures 1B–1C and S1–S3). In all structures, calmodulin (CaM) binds to KCNQ1 at a 1:1 stoichiometry, as an obligate structural component^28-30,37^ (Figures 1B and S1A; Table S1). Through comprehensive 3D classification of the KCNQ1–KCNE1 (-PIP2) particles, we observed complexes with different, up to 4:4, stoichiometry (Figure S3D), a topic of considerable debate in the field^38-43^. Our complex was purified in a detergent solution, which might not fully represent the native cellular environment; nonetheless, our results support the possibility of a 4:4 stoichiometry^42,43^

Analyses of KCNQ1, KCNQ1–KCNE3, and KCNQ1–KCNE1 structures (Figure S4) reveals similarities in overall architecture. Transmembrane domains of KCNQ1 and KCNQ1–KCNE1/3 exhibit a classic domain-swapped architecture^28,29,44^. Binding of KCNE1/3 induces slight clockwise rotations of the voltage sensor domain viewed from cytosol (Figure S4A), while pore domains are nearly identical with a root mean square deviation (RMSD) less than 0.5 Å. As previously observed in the KCNQ1–KCNE3 complex^29^, PIP2 binding to KCNQ1–KCNE1 promotes a transition in the KCNQ1 S6-HA helices from a bent (-PIP2) to a straight (+PIP2) conformation (Figures 1B–1C and S4B). In contrast, KCNQ1 with PIP2 was captured in both bent and straight conformations (Figure 1B), indicating that KCNE1/3 may facilitate the conformational transition. The bent-to-straight shift correlates with significant pore dilation (Figure S4C–S4D) and is expected to increase potassium permeation, providing a general structural mechanism for the PIP2-dependent activation of KCNQ1 and its KCNE1/3 complexes.

Like KCNE3^29^, KCNE1 tucks its transmembrane helix within a cleft formed by three neighboring KCNQ1 subunits (Figure S5A). In the KCNQ1–KCNE1 complex, the voltage sensors are displaced slightly downward (Figure S5B), and packing between KCNE1 and KCNQ1 is looser compared to KCNQ1–KCNE3 (Figures S5C–S5G). The looser packing aligns with the lower biochemical stability of KCNQ1–KCNE1 relative to KCNQ1–KCNE3. The interactions between KCNE1 and KCNQ1, along with the comparison to KCNE3, are shown in detail in Figures S5B– S5H.

### A second PIP2 binding site in KCNQ1

A notable feature of the KCNQ1–KCNE1 complex is the presence of two PIP2-binding sites per KCNQ1 protomer, adding up to eight PIP2 molecules in the entire channel complex (Figure 2A), whereas only one PIP2 per protomer had been previously reported in the KCNQ1–KCNE3 complex^29^. The first PIP2 is bound near the turn between the S4 helix and the S4-S5 linker (Site 1) (Figure 2A). Site 1 is formed mainly by S4, S2-S3 and S4-S5 linkers of the voltage sensor and by KCNE1 (Figures 2A and S6A–S6D) and is similar to the reported site in KCNQ1– KCNE3^29^. We also observed PIP2 bound to this position in KCNQ1 alone, in both bent and straight conformations (Figures S6A–S6B); in the bent conformation, CaM also interacts with PIP2.

**Figure 2:**
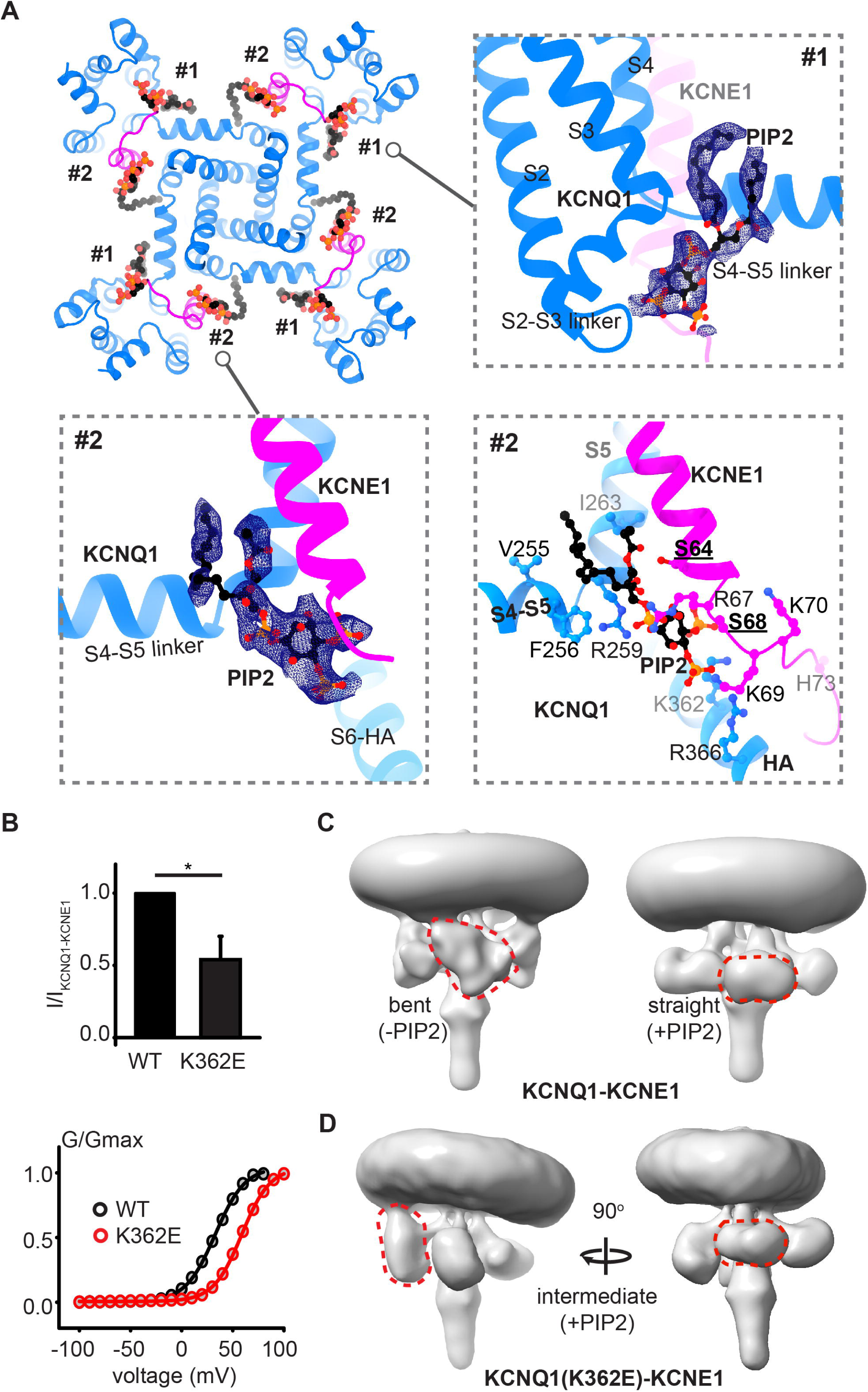
PIP2-binding sites in the KCNQ1–KCNE1 complex. **A**. PIP2-binding sites in the KCNQ1–KCNE1 complex. KCNQ1, KCNE1 and PIP2 are colored in blue, magenta and black, respectively. Cryo-EM density with a ball-and-stick model of PIP2 are shown for both sites. For Site 2, a zoom-in view of the PIP2-binding pocket with key residues are displayed. **B**. Top: electrophysiology characterization of the KCNQ1^K362^ mutation. I/I_KCNQ1–KCNE1_ of wild type (wt) and K362E mutation at 40mV is plotted as bar graphs (n>10, **: p ≤ 0.01). Bottom: G-V curve of the wt (black) and K362E (red) mutation. V_1/2_ of wt and K362E are 31.42 mV and 57.18 mV, respectively. **C**. The conformational changes in the cytosolic domain of KCNQ1 from bent to straight conformation upon PIP2 activation. The orientation changes of cytosolic domain of KCNQ1 and CaM are indicated by red dashed circles. **D**. The conformational changes in the cytosolic domain of KCNQ1^K362E^–KCNE1 mutation in the presence of PIP2. The cytosolic domains present different orientations within one complex.

The second PIP2-binding site in KCNQ1–KCNE1 is formed by the S4-S5 linker, S5, S6-HA helices and KCNE1 (Site 2) (Figures 2A and S6E–S6H). The high-resolution structure of KCNQ1–KCNE1 with PIP2 enables a detailed examination of Site 2 (Figure 2A). The positively charged binding pocket (Figure S6F) includes residues Val255, Phe256, and Arg259 from the S4-S5 linker; Ile263 from S5; Lys362 and Arg366 from the S6-HA helix; and Ser64, Arg67, Ser68, Lys69, Lys70 and His73 from KCNE1. Many of the KCNQ1 residues are critical for coupling the voltage sensor and pore domains^26^.

To explore the significance of PIP2-binding Site 2, we conducted mutagenesis followed by electrophysiology and structural characterization. Mutations of key polar or positively charged residues at Site 2 affect channel gating and/or current amplitude (Figures S7A–S7F). We also determined the cryo-EM structure of the KCNQ1^K362E^ mutant with KCNE1, which shows reduced current amplitude and right-shifted voltage-dependent activation (Figure 2B) and found that this mutation prevented the formation of a stable straight conformation, trapping the channel in intermediate states (Figures 2C–2D). A similar effect was observed with the KCNQ1^R259Q^– KCNE1 mutant complex (Figures S7F–S7G).

Our findings of the PIP2-binding Site 2 are consistent with previous work showing that mutations near Site 2 significantly reduce KCNQ1-mediated currents without affecting voltage sensor movement, regardless of KCNE1 presence^26^. Moreover, KCNE1 was shown to increase the PIP2 binding affinity via basic residues from the cytosolic side of its transmembrane helix, including Arg67, Lys69, and Lys70^33^, which contribute to PIP2-binding Site 2 in the structure (Figure 2A). Together, we conclude that PIP2 binding at Site 2 is crucial for the conformational transition to, and stabilization of, the straight state, which is required to facilitate the coupling between the voltage sensor and pore domains for channel opening.

Our findings on PIP2-binding Site 2 in KCNQ1–KCNE1 prompted us to reevaluate the presence of this site in KCNQ1 and KCNQ1–KCNE3^29^. We first attempted to model PIP2 at Site 2 into the cryo-EM densities (Figures 3A and S6E–S6F). By setting the contour level of cryo-EM densities to the same level as Site 1, we identified potential PIP2 densities in activated KCNQ1–KCNE3 (EMD-20967)^29^, which had been previously overlooked, and in the straight conformation of KCNQ1. The bent conformation of KCNQ1 (+PIP2) lacks density at Site 2, suggesting that PIP2 binding at Site 1 alone is insufficient for KCNQ1 activation (Figures S4C–S4D). The interactions between PIP2 at Site 2 and KCNQ1 or KCNQ1–KCNE3 are detailed and analyzed in Figures S6E–S6F. Taken together, we conclude that each KCNQ1 or KCNQ1–KCNE1/3 protomer accommodates two PIP2 molecules in the straight conformation.

**Figure 3:**
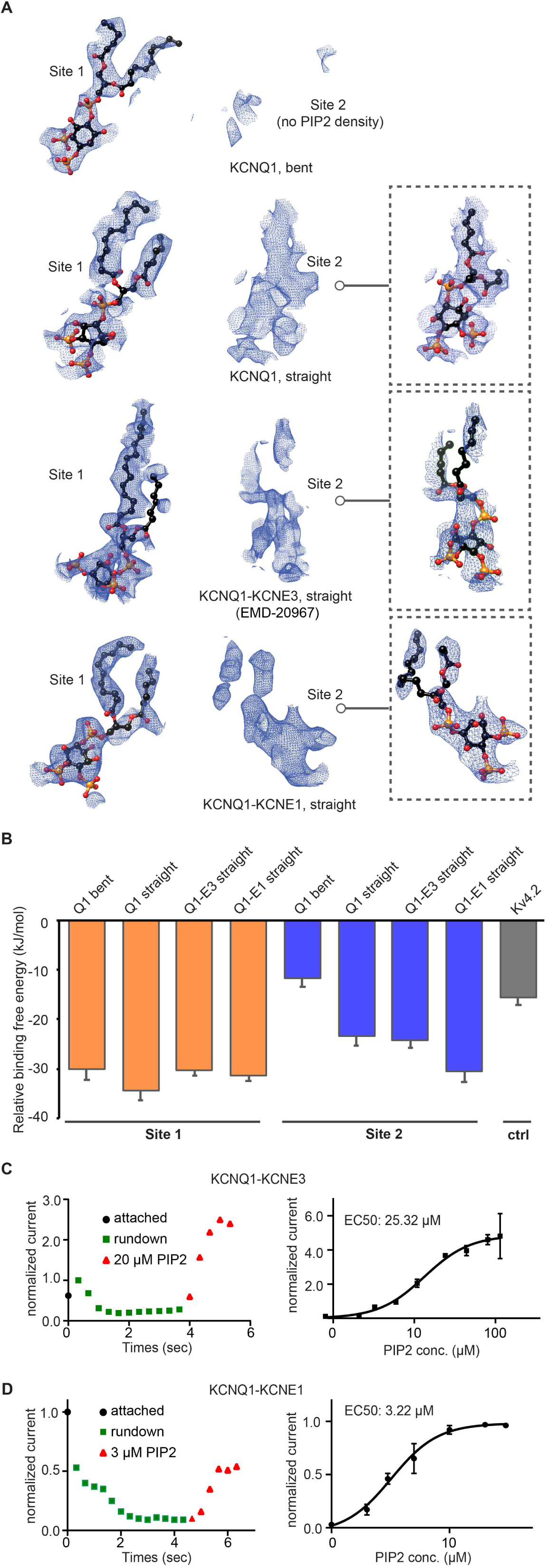
PIP2-binding sites in KCNQ1 and KCNQ1–KCNE complexes. **A**. Cryo-EM density of two PIP2-binding sites in KCNQ1 bent, KCNQ1 straight, KCNQ1–KCNE3 straight, and KCNQ1–KCNE1 straight conformations. The two sites in each complex are contoured at the same level. PIP2 molecules are modeled into Site 2 in the insets. **B.** CG-FEP simulations results of the relative binding free energy of PIP2 to each binding site for the complexes in **A**. We use the binding of Kv4.2 to PIP2 as a negative control for non-specific binding, which has not been reported to bind PIP2. (mean ± SEM, n=15). n stands for number of FEP repeats. **C–D.** Measurement of KCNQ1–KCNEs recovery from current rundown by PIP2 titration and their EC50. **Left**: the represent current recovery trace at the PIP2 concentration near EC50. **Right**: curve fitting of the titration experiment with EC50 indicated. The Hill coefficient was 1.35 and 2.21, respectively, for the KCNE3 and KCNE1 traces. Data are presented as mean ± SEM, n ≥3.

### Validation of the PIP2-binding sites

Coarse-grained Free Energy Perturbation (FEP) and MD simulations were conducted to validate and estimate the binding affinity of PIP2 in both bent and straight conformations of KCNQ1, as well as for KCNQ1–KCNE3 and KCNQ1–KCNE1 in the straight conformation (Figures 3B and S8A). In agreement with structural observations, PIP2 binds to KCNQ1 and KCNQ1–KCNE1/3 at Site 2 in the straight conformation, with varying levels of free binding energies (Figure 3B). In contrast, KCNQ1 in the bent state showed significantly lower binding free energy, comparable to the negative control Kv4.2, which is consistent with the absence of PIP2 density at Site 2 (Figure 3A). Simulations of PIP2 binding at Site 1 showed similar binding free energies across all complexes and conformations (Figure 3B). The binding free energies, indicative of the theoretical PIP2-binding affinities, suggests that Site 1 has a high affinity to PIP2, and that KCNQ1–KCNE1 has a higher PIP2-binding affinity than KCNQ1 alone or KCNQ1–KCNE3 at Site 2. This led us to speculate that the association with different KCNEs results in varied PIP2-binding affinities and sensitivities in KCNQ1.

To test the hypothesis and support computational predications, we monitored the PIP2 dose-dependent recovery of KCNQ1–KCNE3 and KCNQ1–KCNE1 channels from current rundown, results of which were believed to be indicators of PIP2-binding affinities^33^ (Figures 3C–3D). KCNQ1 and KCNQ1–KCNE1/3 complexes exhibit current rundown in excised membrane patches^33^, likely due to PIP2 degradation or diffusion out of the membrane, and external PIP2 application could restore the potassium current in a dose-dependent manner. We found that KCNQ1–KCNE1 recovered more readily than KCNQ1–KCNE3, with apparent EC50 values of 3.22 μM and 25.32 μM (Figures 3C–3D), respectively. Furthermore, adding excess PIP2 to excised membrane patches increased the maximum current of KCNQ1–KCNE3 channels beyond their baseline levels but not of KCNQ1–KCNE1 channels (Figures 3C–3D). These data suggest that the endogenous PIP2 concentration in the oocyte membrane is adequate to activate KCNQ1–KCNE1 channels but insufficient to fully open KCNQ1–KCNE3 channels. Combined with our previous findings that KCNE1 enhances the same EC50 of KCNQ1 by over 100-fold^33^, we conclude that KCNE1 and KCNE3 association lead to varying PIP2 sensitivities, ranked as: KCNQ1–KCNE1 > KCNQ1–KCNE3 > KCNQ1 alone.

MD simulations indicate that residues like S64 and S68 in KCNE1 have higher contact frequencies than their counterparts, G78 and S82 in KCNE3 (Figure S8B). Structurally, KCNE1 appears to displace the PIP2 molecule at Site 2, leading to shorter distances between the PIP2 head group and the positive residues (K362 and R362) on the S6-HA helix (Figures S6E and S6G), contacts with R539 of KCNQ1 that are absent in KCNQ1–KCNE3 complexes (Figures S6G and S8D). These structural differences provided a plausible explanation to the observed higher PIP2 affinity and sensitivity at Site 2 of KCNQ1–KCNE1.

### Sensitivity to Gq-coupled GPCR signaling

PIP2 is degraded by PLC upon Gq-coupled GPCR activation. Thus, in light of their different PIP2 affinities and sensitivities, KCNQ1–KCNE1 or KCNQ1-KCNE3 complexes should display diverse responses to GqPCR signaling (Figure 4A). To test this hypothesis, we co-expressed Gαq-coupled GPCR with KCNQ1 or KCNQ1–KCNE1/3 complexes in *Xenopus* oocytes. We used M1R (muscarinic acetylcholine receptor M1), a predominantly Gαq protein-coupled receptor^45^, and measured channel activities before and after the application of M1R agonist Oxotremorine-M (Oxo-M)^46^ (Figures 4A–4B and S9). Activation of M1R by Oxo-M reduced the potassium currents of KCNQ1, KCNQ1–KCNE3, and KCNQ1–KCNE1 to approximately 6%, 18%, and 76%, respectively, of initial current amplitudes. In fact, we observed significant current reduction for KCNQ1 and KCNQ1–KCNE3, but not for KCNQ1–KCNE1, in response to the basal activity^47^ of M1R, without Oxo-M application (Figure 4B). These findings suggest that the higher sensitivity of KCNQ1 and KCNQ1-KCNE channel complexes to Gq-coupled GPCR signaling correlates with their lower apparent PIP2 affinity, observed in excised membrane patches and MD simulations.

**Figure 4:**
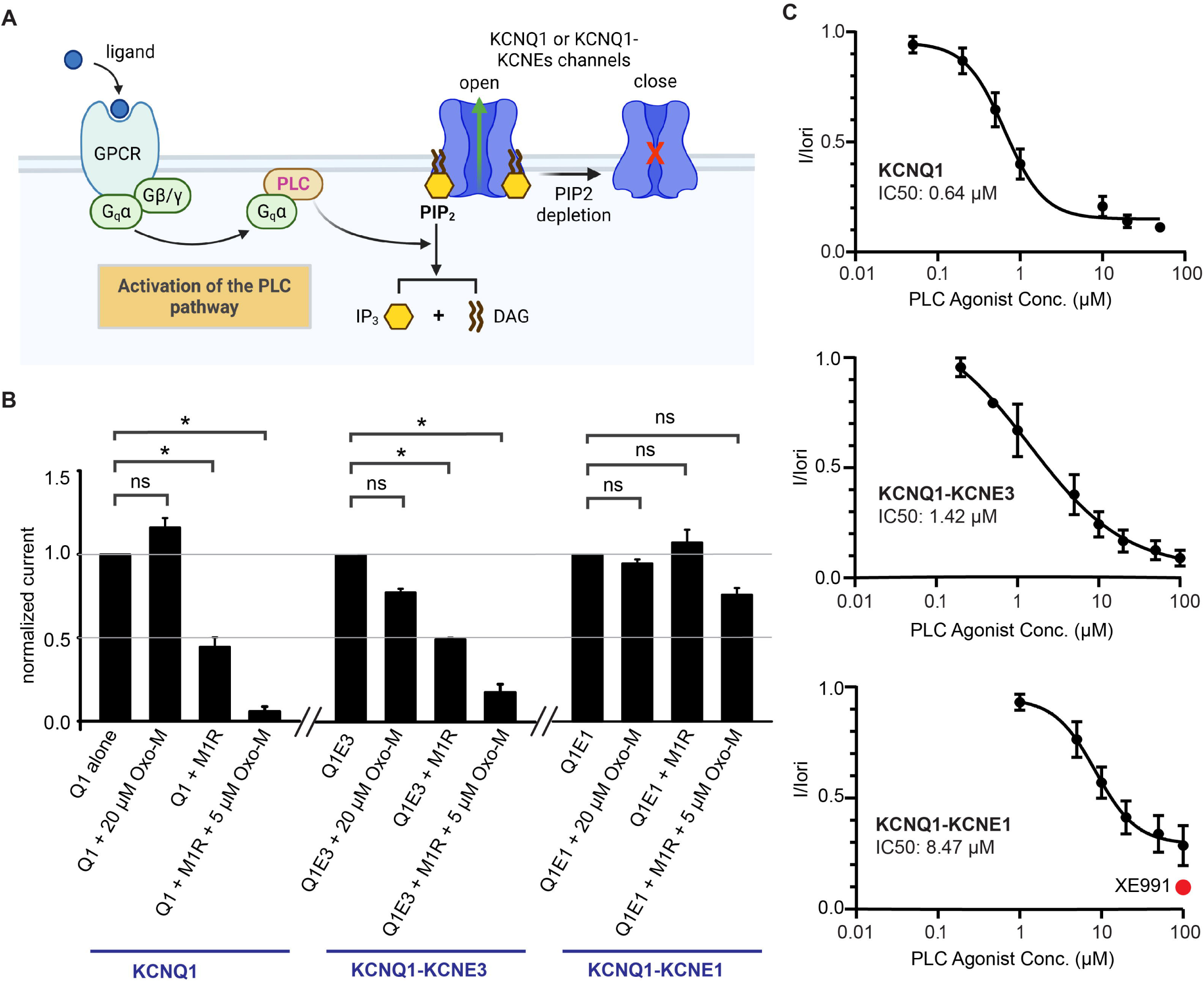
Sensitivity of KCNQ1 and KCNQ1–KCNE complexes to GqPCR pathway. **A**. A cartoon showing the pathway that we reconstituted in Oocytes to test the sensitivity of KCNQ1 and KCNQ1–KCNEs to GqPCR pathway. **B**. Normalized current of KCNQ1 and KCNQ1–KCNE complexes in response to M1R activation by Oxo-M. Q1 alone, Q1E3, and Q1E1 stands for KCNQ1, KCNQ1–KCNE3 and KCNQ1–KCNE1 complexes, respectively. M1R represents the Gq-coupled muscarinic acetylcholine receptor M1. Oxo-M is an agonist of M1R. *ns*: no significant differences; *\*\*\*\*:* p ≤ 0.0001, n is the number of oocytes recorded. For KCNQ1 and KCNQ1–KCNE1 groups, n ≥ 10. For the KCNQ1–KCNE3 group, n ≥ 6; **C**. Curve fitting of the PLC agonist titration experiments in Figure S10 with IC50 indicated. PLC agonist failed to inhibit the KCNQ1–KCNE1 completely with 100 mM agonist. A complete inhibition of the KCNQ1–KCNE1 was indicated using a KCNQ1 inhibitor, XE991 (red data dot). Data are presented as mean ± SEM, n ≥ 5. Hill co-efficient of the curves are -1.814 for KCNQ1, -0.7389 for KCNQ1–KCNE3, and -1.724 for KCNQ1–KCNE1.

To quantitatively evaluate the sensitivity of KCNQ1 channel complexes to GPCR signaling, we employed a PLC agonist *m*-3M3FBS^48^ to mimic GqPCR activation and measured the current inhibition of KCNQ1 and KCNQ1–KCNE1/3 channels (Figures 4C and S10). The IC50 values in response to *m*-3M3FBS were determined to be 0.64 µM for KCNQ1, 1.42 µM for KCNQ1– KCNE3, and 8.47 µM for KCNQ1–KCNE1. Notably, at the highest concentration of the PLC agonist tested (100 µM), the KCNQ1–KCNE1 current was not completely inhibited, as confirmed by application of the KCNQ1 inhibitor XE991 (Figure 4C). This implies that a portion of the *I_Ks_*current, conducted by KCNQ1–KCNE1 complexes in cardiac cells, could be resistant to GqPCR regulation under physiological conditions.

## DISCUSSION

In this study, we analyzed cryo-EM structures of KCNQ1, and KCNQ1–KCNE1/3 complexes with and without PIP2, which revealed two distinct PIP2-binding sites within each KCNQ1 protomer (Figures 1 and 2A). Site 1 was shown to be regulated by membrane potential^49^. Site 2 is believed to be important for coupling voltage sensor and pore domains^26^ and occupied when KCNQ1 is in the straight conformation (Figure 3A). PIP2 binding at Site 2 is crucial for the transition to and stabilization of the straight conformation (Figure 2C), and thus essential for channel activation and ion conductance. Combining structural biology with MD simulations and functional assays (Figures 3B–3D and 4), we further demonstrated that KCNE1 and KCNE3, by contributing significantly to the formation of PIP2-binding Site 2, impart different PIP2 and

GPCR sensitivities to KCNQ1. As a result, KCNQ1 operates as a PIP2-gated channel in non-excitable cells with KCNE3 and as a predominantly voltage-gated potassium channel in excitable cells with KCNE1. The distinct modulation of KCNQ1 channels by KCNE1 and KCNE3 offers valuable insights for developing targeted therapies for KCNQ1-related diseases. For cardiac disorders involving KCNQ1–KCNE1 complexes, pharmacological tools specifically targeting KCNE1 would be beneficial. In contrast, for cystic fibrosis associated with KCNQ1– KCNE3 complexes, targeting Gq-coupled GPCR signaling pathways could be also effective as KCNE3-specific compounds.

The PIP2 binding Site 2 is located at the interface between the voltage sensor and pore domains, believed to facilitate their mechanical coupling. The site is formed by the S0-S1 segments of the voltage sensor domain and the S4-S5 linker, S5, and S6 segments of the pore domain from the neighboring subunit (Figure S6F). PIP2 binding at Site 2 mediates the interaction between these domains, inducing significant conformational changes, including the formation of a continuous helix between S6 and HA and pore dilation (Figure S4, Movies S1-2). The accessory subunits KCNE1 or KCNE3 displace the S0-S1 segment of the voltage sensor domain, occupying its position in the PIP2 binding Site 2 (Figure S6F, Movies S3-4). Differences in PIP2 coordination by KCNE1 and KCNE3 (Figures S6G-H, S8D) may account for variations in PIP2 affinity at Site 2; however, these differences do not fully explain the differences in voltage-dependent gating, which is likely a kinetic problem requiring further investigation. KCNE1 and KCNE3 interact with PIP2 via conserved residues on the inner face of their transmembrane helices. Given that all five human KCNE isoforms modulate KCNQ1 and possess conserved PIP2-binding residues (Figure S5H), we speculate that all KCNQ1-KCNE complexes should contain a second PIP2 binding site, with KCNEs interacting with PIP2 in a comparable manner.

We propose a working model to illustrate the molecular mechanism underlying the cell-specific regulation of KCNQ1 by KCNE1 and KCNE3 (Figure 5). Our research shows that KCNE1 and KCNE3 tune the voltage- and lipid-dependent gating sensitivity of KCNQ1. In excitable cells, KCNQ1 associates with KCNE1 to form a channel complex that is voltage-dependent but less responsive to Gq-coupled GPCR signaling. Indeed, it is well-established that KCNE1 modifies the voltage-dependent gating of KCNQ1 by slowing its activation and deactivation kinetics, thereby facilitating the *I_Ks_* current conduction in cardiac tissues^7,8^. However, modulation of *I_Ks_* current by GqPCR, manifesting a biphasic effect in overexpressed systems^50^, is seemingly contradictory and has yet to be clarified in native cardiac systems^27,51-54^. Additionally, the movement of voltage sensor upon depolarization could disrupt PIP2 binding at Site 1^49^, indicating that voltage-dependent gating might be able to override PIP2 regulation in excitable cells. In non-excitable cells, KCNQ1 associates with KCNE3, forming a channel that remains open under physiological membrane potentials^6,55^. The voltage-insensitive KCNQ1–KCNE3 channel complex is more susceptible to regulation by PIP2 degradation and GqPCR signaling. Although the detailed mechanism underlying the differential voltage-dependent gating between KCNE1 and KCNE3 requires further investigation, our structures provide a framework for future functional and structural studies to elucidate these differences. Nevertheless, we provide a novel molecular framework demonstrating how the multifunctional KCNQ1 channel adjusts its voltage and lipid dependences to fulfill diverse physiological roles across different tissues through association with sequence-homologous KCNE subunits.

**Figure 5:**
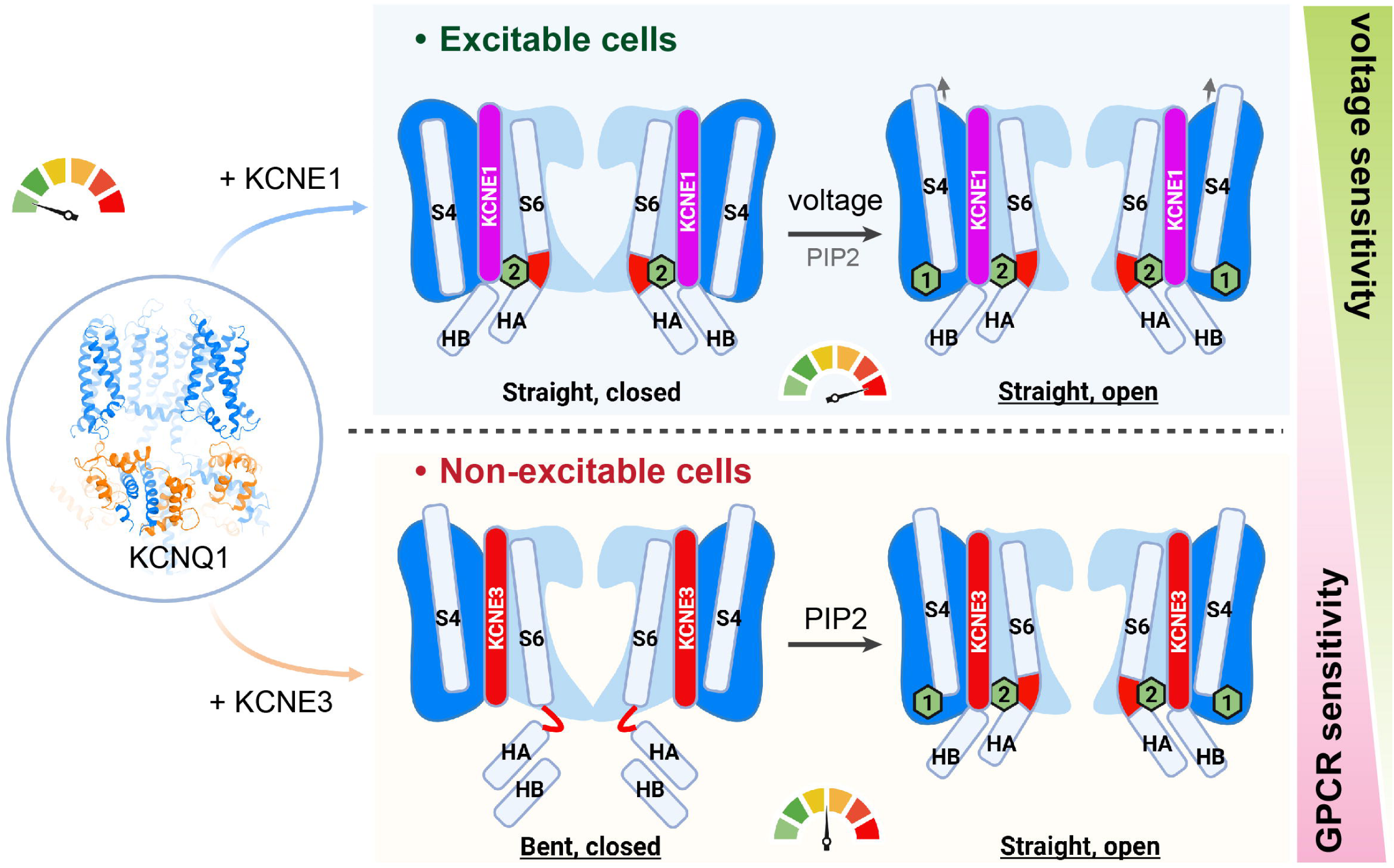
Working model of the cell-specific regulation of KCNQ1. KCNQ1 can form complexes with KCNE1 and KCNE3, and these complexes are activated by voltage and lipid signals with varying sensitivities in a tissue-specific manner. The KCNQ1– KCNE1 complex is predominantly voltage-gated and maintains its function in excitable cells, such as nerve and muscle cells, due to its resistance to PIP2 degradation and GqPCR signaling. Conversely, the KCNQ1–KCNE3 complex is voltage-insensitive and remains constitutively open under physiological conditions, facilitating potassium conduction in non-excitable cells, like epithelial cells, but is more susceptible to GqPCR signaling regulation due to its lower PIP2-binding affinity. PIP2 affinities are qualitatively indicated by color indicators with a pointer for different channels. Cryo-EM structures of the underlined states are available.

KCNE1/3 and PIP2 appear to synergistically regulate KCNQ1, a mechanism that may have broad implications for ion channel modulation by KCNE subunits and PIP2. In KCNQ1, KCNE1/3 facilitate PIP2 binding by contributing to the formation of the PIP2-binding pocket; PIP2, in turn, enhances the interaction between KCNQ1 and KCNE1, as mutations in the PIP2-binding site of KCNQ1 (KCNQ1^R259Q^) result in weaker cryo-EM density for KCNE1 (Figure S7F). These findings suggest a co-modulatory mechanism in which KCNE subunits and PIP2 act as mutual cofactors, each enhancing the other’s ability to regulate ion channel function. This co-modulatory mechanism may extend beyond KCNQ1 to other ion channels regulated by KCNEs and PIP2. For example, the hERG channel is known to be modulated by both KCNE2^56^ and PIP2^57^, but the structural details of this regulation remain poorly understood. The concept of co-modulation could provide a framework to address this gap, offering insights into the molecular basis of such regulatory interactions. Moreover, this mechanism may apply to other ion channels, including HCN, Kv1.2, Kv4, and probably TMEM16A, all of which have been reported to be regulated by KCNE subunits^58-62^ and PIP2^63^. This suggests that the co-modulation mechanism by KCNEs and PIP2 could represent a widespread, yet previously underappreciated, mechanism for fine-tuning ion channel activity.

Together, our work uncovers a novel paradigm of channel modulation whereby cell-specific KCNE beta subunits alter the primary gating signals of KCNQ1, enabling it to fulfill diverse physiological roles across various tissues. These molecular insights provide a deeper mechanistic understanding of the functional plasticity of KCNQ1, highlighting its ability to alter sensitivity between two different gating signals: voltage and lipid. The foundational knowledge establishes a basis for developing strategies to target ion channel regulation in a tissue-specific manner, potentially leading to more precise therapeutic interventions for disorders linked to KCNQ1 dysfunction.

## Supporting information

Supplemental Figure 1, and will be used for the link to the file on the preprint site

Supplemental Figure 2, and will be used for the link to the file on the preprint site

Supplemental Figure 3, and will be used for the link to the file on the preprint site

Supplemental Figure 4, and will be used for the link to the file on the preprint site

Supplemental Figure 5, and will be used for the link to the file on the preprint site

Supplemental Figure 6, and will be used for the link to the file on the preprint site

Supplemental Figure 7, and will be used for the link to the file on the preprint site

Supplemental Figure 8, and will be used for the link to the file on the preprint site

Supplemental Figure 9, and will be used for the link to the file on the preprint site

Supplemental Figure 10, and will be used for the link to the file on the preprint site

Supplemental Movie 1, and will be used for the link to the file on the preprint site

Supplemental Movie 2, and will be used for the link to the file on the preprint site

Supplemental Movie 3, and will be used for the link to the file on the preprint site

Supplemental Movie 4, and will be used for the link to the file on the preprint site

## FIGURE LEGENDS

**Figure S1: Structural determination of KCNQ1–KCNE1 in the presence of PIP2**

**A.** Expression, purification and reconstitution of KCNQ1^I145C^–KCNE1^K41C^ complexes in nanodiscs using MSP2N2 as the membrane scaffold. The SDS-PAGE lane of the purified complex is shown. **B.** Representative electrophysiological recordings of KCNQ1–KCNE1 wt (black) and cysteine mutation pair (red) in different reducing (green) and oxidizing (blue) conditions. The voltage step applied from -100 mV to + 80 mV at 10-mV increments. Holding potential is -80 mV and tail current record at - 40 mV. **C.** The G-V curves with the same color code of the recordings in panel (**B**). The V_1/2_ of each condition is indicated. **D.** The current amplitude of the recordings in panel (**B**) at 40 mV. Data are presented as mean ± SEM, n ≥ 5. ***: p ≤ 0.001; ****: p ≤ 0.0001; ns: not significant. **E.** A brief flow chart of the structure determination process. The local resolution map, angular distribution and resolution FSC at 0.143 curves are shown. **F.** Local cryo-EM density of PIP2 molecules and key secondary structures of KCNQ1 and KCNE1. **G.** Model-to-Map FSC of the KCNQ1–KCNE1 complex. **H.** Modeling of I145C of KCNQ1 and K41C of KCNE1 with nearby cryo-EM density shown.

**Figure S2: Structural determination of KCNQ1 in the presence of PIP2**

**A.** A brief flow chart of the cryoEM structure determination procedure. The local resolution maps are also shown. **B–C.** Angular distribution (left) and resolution Fourier shell correlation (FSC) (right) at 0.143 of the straight conformation (**B**) and bent conformation (**C**) reconstructions. **D.** Model-to-Map FSC of the straight (left) and bent (right) conformations. **E–F.** Cryo-EM density of key secondary structures of KCNQ1 and PIP2 molecules in straight (**E**) and bent (**F**) conformations, respectively.

**Figure S3: Structural determination of KCNQ1–KCNE1 in the absence of PIP2**

**A.** A brief flow chart of the structural determination protocol for KCNQ1–KCNE1 complexes in nanodiscs without PIP2. **B.** Cryo-EM density of key secondary structures of KCNQ1 and KCNE1.

**B.** The FSC curve between cryo-EM map and structural model. **D.** 3D classes and maps of KCNQ1:KCNE1 in different stoichiometry (from 4:4 to 4:1). The cartoon models of KCNQ1 and KCNE1 are colored in blue and magenta, respectively. The cryo-EM density is colored in transparent blue.

**Figure S4: Structural comparison between KCNQ1 and KCNQ1–KCNE1/3**

**A.** The rotation of voltage sensor domains introduced by binding of KCNE subunits viewed from the cytosol side. KCNQ1, KCNQ1–KCNE3, and KCNQ1–KCNE1 are colored in grey, blue and magenta, respectively. Structures are aligned to the pore domains of KCNQ1. **B.** PIP2-caused bent-to-straight transition of KCNQ1 and KCNQ1–KCNE1/3 complexes. **C–D.** The ion conductance path (**C**) and pore radius (**D**) plotted using the HOLE program.

**Figure S5: Structural comparison between KCNQ1–KCNE1 and KCNQ1–KCNE3**

**A.** Binding pocket of KCNE1 (magenta) and KCNE3 (red) in KCNQ1 in both bent and straight conformations. **B.** Alignment of KCNQ1–KCNE1 (blue and magenta) and KCNQ1–KCNE3 (grey) by S4-S5 linker and S5 to show the relative displacement of the pore domain upon KCNE binding. **C–G.** Detailed interactions between KCNQ1 and KCNE1/3 side by side. Three neighboring KCNQ1 subunits are colored in cyan, green and pink, respectively. KCNE1 and KCNE3 are colored in magenta and red, respectively. PIP2 binding sites are indicated by dashed circles. **H.** Sequence alignment between KCNEs. The transmembrane helix is highlighted in yellow. Key residues involving in PIP2 site 2 are labelled with red dots.

**Figure S6: PIP2-binding sites in KCNQ1 and KCNQ1–KCNE complexes**

**A–D.** PIP2-binding pockets at Site 1 in KCNQ1 (+PIP2, bent) (**A**), in KCNQ1 (+PIP2, straight) (**B**), KCNQ1–KCNE3 (+PIP2, straight) (**C**), and KCNQ1–KCNE1 (+PIP2, straight) (**D**). **E.** PIP2-binding pockets at Site 2 (from left to right) in KCNQ4 (PDB 7VNP)^64^, KCNQ1 (+PIP2, straight), KCNQ1–KCNE3 (+PIP2, straight), and KCNQ1–KCNE1 (+PIP2, straight). Here KCNQ4 was used for comparison, for its structure revealed two similar PIP2 binding sites. **F.** Atomic details of the PIP2-binding site 2 in KCNQ1 (+PIP2, straight), KCNQ1–KCNE3 (+PIP2, straight), and KCNQ1–KCNE1 (+PIP2, straight) from left to right. Inset: electrostatic surface representation of the binding pocket. Key residues show different contact frequency in KCNEs based on MD simulations are underlined with bold fonts. **G.** Structural alignment of PIP2-binding Site 2 between KCNQ1–KCNE1 (blue, magenta and black) and KCNQ1–KCNE3 (grey). The shortest distances between PIP2 and C-alpha of K362 and R366, and between PIP2 and D537 and I542 are shown. D537 and I542 are the two resolved residues to indicate the distance between PIP2 and R539, which was not resolved in our cryo-EM structure. **H.** Comparison between KCNQ1, KCNQ1, KCNQ1–KCNE1 and KCNQ1–KCNE3, which shows the different voltage sensor displacement upon PIP2 binding at Site 2.

**Figure S7: Structural and functional characterization of mutations at Site 2**

**A–D.** Current amplitude and G-V curve of Site 2 mutations with or without KCNE1. **A**: KCNQ1 R259Q/E; **B**: KCNQ1 K362Q/E; **C**: KCNQ1 R366Q/E; **D**: KCNE1 R67Q, K69Q, R67Q+K69Q, and S68E. The bar graphs are plotted using currents recorded at 40 mV. Statistical significances are annotated as follows: ns: p > 0.05, not significant; *: p ≤ 0.05; **: p ≤ 0.01; ***: p ≤ 0.001; ****: p ≤ 0.0001. For all experiments in Figures S7A-D, n ≥ 10. **E.** Current amplitude changes in Site 2 when KCNE3 mutations are introduced at corresponding residues as KCNE1. Same statistical significance annotation is used as in Figures S7A-D, n=8. **F.** Structural determination of KCNQ1^R259Q^–KCNE1 complexes in PIP2 nanodiscs. Similar procedure was used for KCNQ1^K362E^–KCNE1. The structures indicated by colored hexagons are used for comparison in structural comparison in (**G**). **G.** Structure comparison between the KCNQ1^K362E^– KCNE1 mutation and the wild type. The cytosolic domain orientation is indicated by a red dashed circle. Structures are filtered to 20 Å for comparison.

**Figure S8: MD simulation of PIP2 binding**

**A.** RMSD of PIP2 in Sites 1 and 2, indicating the stability of the PIP2 molecule in its binding site.

**B.** Contact frequency of key residues in KCNE1 and KCNE3 to the PIP2 molecule bound in Site

**C–D**. Contact frequency of key residues in KCNQ1 to the PIP2 molecule bound Sites 1 and 2. Data are plotted as mean ± SEM, n = 12 (4 PIP2 molecules for each channel with 3 repeats).

**Figure S9: GPCR activation and KCNQ1–KCNE1 current**

**A.** Raw traces of KCNQ1–KCNE1 in response to different concentrations of Oxo-M treatment. Traces are colored with the same color codes used in (**B**) and (**C**). For KCNQ1–KCNE1, raw traces recorded from -100 mV to +80 mV at 10-mV increments, for KCNQ1 and KCNQ1– KCNE3, raw traces recorded from -100 mV to +60 mV at 10-mV increments. **B**. The current traces at 40 mV upon application of different concentrations of Oxo-M. **C.** The G-V curve of KCNQ1–KCNE1 in the presence of different concentrations of Oxo-M. The V_1/2_ are indicated. Data are plotted as mean ± SEM, n >3.

**Figure S10: Representative traces of PLC titration and channel inhibition**

**A.** Representative raw traces of KCNQ1–KCNE1 channel activity in response to varying concentrations of m-3M3FBS. Traces are color-coded, with consistent color usage in panels **B** and **C**. Voltage steps were applied from -100 mV to 80 mV in 10 mV increments. **B.** Conductance-voltage (G-V) curve for KCNQ1–KCNE1 in the presence of different *m*-3M3FBS concentrations. Data are presented as mean ± SEM, with n >6. **C.** Current traces recorded at 40 mV with the application of various *m-*3M3FBS concentrations. **D.** Raw traces depicting KCNQ1– KCNE3 channel responses to different *m*-3M3FBS concentrations. Traces are color-coded, with the same colors used in panel **E.** Voltage steps were applied from -100 mV to 60 mV in 10 mV increments. **E.** Current traces at 40 mV with the application of *m-*3M3FBS at different concentrations. **F.** Representative raw traces of KCNQ1 channels alone under varying concentrations of m-3M3FBS. Consistent color-coding is maintained in panels g and h. Voltage steps were applied from -100 mV to 60 mV in 10 mV increments. **G.** G-V curve for KCNQ1 in the presence of different m-3M3FBS concentrations. Data are presented as mean ± SEM, with n > 6 experiments. **H.** Current traces recorded at 40 mV upon application of varying concentrations of *m*-3M3FBS.

**Movie S1: Site 2 PIP2-induced conformational change of KCNQ1 at the protomer level**

**Movie S2: Site 2 PIP2-induced conformational change of KCNQ1 at the pore region**

**Movie S3: PIP2-binding site remodeling by KCNE1**

**Movie S4: PIP2-binding site remodeling by KCNE3**

**Table S1:** Cryo-EM data collection, refinement and validation statistics.

|  | KCNQ1–KCNE1<br>(PIP2-free)<br>(EMDB-XXXX)<br>(PDB XXXX) | KCNQ1–KCNE1<br>(+PIP2)<br>(EMDB-XXXX)<br>(PDB XXXX) | KCNQ1 (+PIP2) |  |
| --- | --- | --- | --- | --- |
|  |  |  | Bent<br>(EMDB-XXXX)<br>(PDB XXXX) | Straight<br>(EMDB-XXXX)<br>(PDB XXXX) |
| **Data collection and processing** |  |  |  |  |
| Magnification | 130,000 | 130,000 | 130,000 |  |
| Voltage (kV) | 300 | 300 | 300 |  |
| Electron exposure ($e^{-}/\text{\AA}^2$) | 51.18 | 52.47 | 52 | |
| Defocus range ($\mu\text{m}$) | 0.5–2.0 | 0.5–2.0 | 0.6–1.8 | |
| Pixel size ($\text{\AA}$) | 0.6485 | 0.6485 | 0.6485 | |
| Symmetry imposed | C4 | C4 | C4 |  |
| Initial particle images (no.) | 2,190,512 | 1,525,191 | 1,984,244 | 1,984,244 |
| Final particle images (no.) | 148,708 | 110,909 | 93,105 | 283,225 |
| Map resolution ($\text{\AA}$) | 2.7 | 3.9 | 3.8 | 3.7 |
| FSC threshold | 0.143 | 0.143 | 0.143 | 0.143 |
| Map resolution range ($\text{\AA}$) | 2.5–30 | 2.5–30 | 2.5–30 | 2.5–30 |
| **Refinement** |  |  |  |  |
| Initial model used<br>(PDB code) | 6v00 | 6v01 | 6uzz | 6v01 |
| Model resolution ($\text{\AA}$) | 3.05 | 4.05 | 3.93 | 3.78 |
| FSC threshold | 0.5 | 0.5 | 0.5 | 0.5 |
| Model resolution range ($\text{\AA}$) | 2.5–30 | 2.5–30 | 2.5–30 | 2.5–30 |
| Map sharpening $B$ factor ($\text{\AA}^2$) | -108 | -203 | -145 | -174 |
| Model composition |  |  |  |  |
| Non-hydrogen atoms | 31160 | 16180 | 29180 | 13228 |
| Protein residues | 2088 | 2200 | 1984 | 1900 |
| Ligands | CA: 8 | PT5: 8, CA: 8 | PT5: 4, CA: 8 | PT5: 8, CA: 8 |
| $B$ factors ($\text{\AA}^2$) | | | | |
| Protein | 55.42 | 55.95 | 31.16 | 26.28 |
| Ligand | 63.50 | 86.06 | 29.15 | 19.78 |
| R.m.s. deviations |  |  |  |  |
| Bond lengths ($\text{\AA}$) | 0.01 | 0.003 | 0.004 | 0.003 |
| Bond angles ($^{\circ}$) | 0.659 | 0.585 | 0.538 | 0.516 |
| **Validation** |  |  |  |  |
| MolProbity score | 1.48 | 1.60 | 1.34 | 1.36 |
| Clashscore | 5.74 | 9.16 | 3.91 | 5.98 |
| Poor rotamers (%) | 0.00 | 0.00 | 0.00 | 0.00 |
| Ramachandran plot |  |  |  |  |
| Favored (%) | 97.07 | 97.43 | 97.08 | 97.85 |
| Allowed (%) | 2.93 | 2.57 | 2.92 | 2.15 |
| Disallowed (%) | 0.00 | 0.00 | 0.00 | 0.00 |

## DATA AVAILABILITY

### Material availability

Plasmids generated in this study are available upon request.

### Data and code availability

All tpr files associated with MD simulations in this study are available through this link at Zenodo (https://zenodo.org/records/14632060). The cryo-EM maps of KCNQ1(+PIP2) and KCNQ1– KCNE1 (+/-PIP2) have been deposited in the Electron Microscopy Data Bank under the accession codes EMD-xxxxx (KCNQ1 with PIP2, bent), EMD-xxxxx (KCNQ1 with PIP2, straight), EMD-xxxxx (KCNQ1–KCNE1, bent) and EMD-xxxxx (KCNQ1–KCNE1 with PIP2, straight). The corresponding coordinates have been deposited in the Protein Data Bank under the accession codes xxxx, xxxx, xxxx and xxxx.

## ACKNOWLEDGMENTS

We thank the staff at the cryo-EM facility at St Jude Children’s Research Hospital for help with data collection, Ines Chen for constructive feedback on manuscript writing and members from Sun lab, including Xiao Chen, Patricia Hixson and Hanwen Zhu for helpful discussions and support. This research is funded by American Lebanese Syrian Associated Charities (ALSAC), President Young Professorship (PYP) from National University of Singapore, and NIH R00HL143037 (to J. Sun); US - Israel BSF research grant 2019159, NIH R01 HL155398 and R01 HL166628 (to J. Cui); and the Knut and Alice Wallenberg Foundation, the Science for Life Laboratory, the Swedish eScience Research Center and Swedish Research Council grants VR 2019-02433 and 2022-04305 (to L. Delemotte). The National Academic Infrastructure for Supercomputing in Sweden (NAISS) and the Swedish Research Council through grant agreement no. 2022-06725 (to L. Delemotte) funded the MD simulations.

## AUTHOR CONTRIBUTIONS

J. Sun and J. Cui conceived, designed and supervised the study. A. Kermani and S. Chittori collected cryo-EM data. C. Cui and A. Kermani performed cryo-EM data process and analyzed the structures under the supervision of J. Sun, C. Cui, A. Kermani and M. Jiang did model building. L. Zhao. and S. Du. conducted the electrophysical experiments and related data analysis under the supervision of J. Cui. T. Pipatpolkai performed the MD simulations and analysis under supervision of L. Delemotte. Y.Z. Tan and J. Shi provided intellectual and technical support during the project. J. Sun prepared the manuscript draft with input from all authors.

## DECLARATIONS OF INTERESTS

The authors declare no conflicts of interest.

## MATERIALS AND METHODS

### Cell lines

Sf9 cells were cultured in Sf-900 II SFM medium (GIBCO) at 27°C. HEK293S GnTI− cells were cultured in Freestyle 293 medium supplemented with 2% fetal bovine serum at 37°C.

### Protein expression and purification

Construct design, protein expression and purification, nanodisc reconstitution for KCNQ1 and KCNQ1–KCNE1 complexes followed the same protocol as the previous KCNQ1–KCNE3 study^29^, with KCNE1 replacing the KCNE3. Briefly, N- and C-terminus loop of KCNQ1 are truncated for stability, resulting a construct including residues 76-620. For protein purification and monitoring, KCNQ1 is tagged with C-terminal GFP and KCNE1 is tagged with N-terminal mCherry tag. I145C and K41C mutations were introduced to KCNQ1 and KCNE1, respectively, for KCNQ1–KCNE1 stabilization and purification. The KCNQ1 and KCNE1 (Homo sapiens: NP_000210.2) coding cDNA were synthesized by Genewiz, Azenta Life Sciences.

### Cryo-EM sample preparation and data collection

Cryo-EM grids were prepared using a Vitrobot Mark IV (FEI). Quantifoil R1.2/1.3 holey carbon gold grids (400 mesh) were glow-discharged for 30 seconds. The concentrated protein sample was mixed with 120 mM Fos-Choline-8 at a volume ratio of 1:50 immediately before applying to the grid and vitrification. In the case of KCNQ1–KCNE1 (+/-PIP2), 3.5 μL of ∼3.3 mg/mL protein sample was applied to each grid, which were double-blotted for 5 seconds under blot force 0 at 100% humidity and 16 °C, then vitrified by plunging into liquid ethane cooled by liquid nitrogen. For KCNQ1 (+PIP2) sample preparation, 3.5 μL of ∼3.5 mg/mL protein sample was applied to a same vitrification protocol as KCNQ1–KCNE1 (+/-PIP2).

All datasets were acquired on 300 keV Titan Krios microscope (FEI) equipped with a K3 direct electron detector (Gatan) using EPU software with magnification of 130,000. Data collection was conducted in super-resolution mode with pixel size of 0.6485 Å. For data collection of KCNQ1-KCNE1 (-/+PIP2) and KCNQ1 (+PIP2), images were recorded with a defocus range of −0.5 to −2.0 μm. Data were collected at a dose rate of ∼10.8 to ∼15.8 e/frame/s with images captured over 50 frames.

### Image processing and 3D reconstruction

Image stacks were gain-normalized and corrected for beam-induced motion using MotionCor2^65^. The contrast transfer function parameters were estimated from motion-corrected summed images without dose-weighting using CTFFIND4^66^. All subsequent processing steps were performed on motion-corrected, dose-weighted summed images. The data process was carried out in CryoSPARC (version 4.6.2)^67^ and RELION^68^.

KCNQ1KCNE1 (-PIP2) data processing was performed in CryoSPARC. 2D classification was conducted to remove junk particles. ∼715k particles were sorted out through *Ab-Initio* Reconstruction and Heterogeneous Refinement with C1 symmetry. Non-uniform refinement generated a map for KCNQ1–KCNE1 with C4 symmetry. Due to the various stoichiometry of KCNQ1 and KCNE1, heterogeneity limits the local resolution of KCNE1. To solve this issue, KCNQ1–KCNE1 map was processed in ChimeraX^69^ with volume eraser to remove KCNE1. Finally, various stoichiometry of KCNE1 to KCNQ1 maps were produced and used as initial volumes. Another round of Heterogeneous Refinement successfully sorted out particles with KCNQ1-KCNE1 stoichiometry of 4:4. Non-uniform refinement was performed with C4 symmetry to generate the final map.

Data processing of KCNQ1–KCNE1(+PIP2) was similar with KCNQ1–KCNE1(-PIP2), while KCNQ1–KCNE1(+PIP2) particles of stoichiometry of 4:4 were sorted out through 3D classification in CryoSPARC. Then particles were extracted to RELION through pyem^70^. ∼111k particles were subjected to 3D auto-refine with C4 symmetry. A mask was generated in RELION excluding HD helix and micelle and input to local 3D auto-refine. After postprocess, map was generated with an overall resolution of ∼3.9 Å.

For data process of KCNQ1 (+PIP2), good particles were classed out after 2D classification. 6 initial volumes were generated from *Ab-Initio* Reconstruction and were used as an input for Heterogeneous Refinement to sort out particles with different conformations. KCNQ1 (+PIP2) bent and straight conformations were confirmed and refined to near-atomic resolution.

### Structural refinement and model building

Models were built in Coot^71^. Initially, KCNQ1-KCNE3 bent (PDB 6V00), KCNQ1-KCNE3 straight (PDB 6V01), KCNQ1 bent (PDB 6UZZ) and KCNQ1 of KCNQ1-KCNE3 straight (PDB 6V01) were docked into the cryo-EM maps of KCNQ1-KCNE1 bent, KCNQ1-KCNE1 (+PIP2) straight, KCNQ1 (+PIP2) bent and KCNQ1 (+PIP2) straight, respectively. Manual rebuilding was then conducted using Coot. The KCNE3 main chain was used as an initial model for building KCNE1 into KCNQ1-KCNE1 electron density map. PIP2 molecules were generated as CIF files by the phenix.eLBOW^72^ and imported as PT5 during model building in Coot. The structural model was iteratively refined using phenix.real_space_refine^73^ with secondary structure restraints and checked in Coot. The quality of the structures was assessed using the MolProbity server^74^.

The pore radii were calculated using HOLE^75^. Figures were created using PyMOL (The PyMOL Molecular Graphics System, Version 2.6.0 and 3.1.1, Schrödinger, LLC) and UCSF Chimera^76^. The electrostatic representations in all figures were calculated using the PyMOL-APBS Plugin^77^.

### Coarse-grained free energy perturbation (CG-FEP)

The side chains and hydrogen atoms of KCNQ1, KCNQ1–KCNE1, and KCNQ1-KCNE3 in complex with CaM, both in bent and straight conformations, were modeled using SwissModel^78^ based on the cryo-EM structures described in the manuscript. Kv4.2 (PDB 7UK5)^79^ served as a negative control for non-specific binding in our study. The initial PIP2 binding site on Kv4.2 is obtained by aligning the pore domain of Kv4.2 to the pore domain of the KCNQ1 channel. The structural data was transformed from an all-atom representation to a coarse-grained model utilizing martinize.py^80^. Subsequently, the protein was embedded within a 1-palmitoyl-2-oleoyl-sn-glycero-3-phosphocholine (POPC) lipid bilayer and solvated in 0.15 M NaCl with coarse-grained water particles using insane.py^81^. This process yielded a simulation box measuring 13 x 13 x 13 nm^3^, containing approximately 22,000 coarse-grained particles. For all simulations conducted in this study, the Martini 2.3 force field was employed.

Simulations were performed using GROMACS 2021.3^82^, employing the V-rescale^83^ thermostat for temperature coupling and the Parrinello-Rahman barostat^84^ for semi-isotropic pressure control. The system’s temperature was maintained at 323 K, and the pressure kept at 1 bar in the xy plane. In this study, a positional restraint force of 1000 kJ mol^-1^ nm^-2^ was applied to the protein backbone. Energy minimization was conducted using the steepest descent algorithm with an energy threshold of 1000 kJ mol^-1^ nm^-2^. The system was then equilibrated for 50 ns prior to CG-FEP calculation.

In the CG-FEP calculation, the coarse-grained PIP2 molecules were then placed in the binding site based on the cryo-EM structure and equilibrated for a further 500 ns. In the simulation with Kv4.2, the additional 500 ns equilibration was omitted to prevent PIP2 from diffusing away from the binding site. The PIP2 headgroup was alchemically transformed into other PC using an approach similar to the previous study^85^ to assess the role of PIP2 during the conformational transition. This was done in chemical space with λ as a coordinate parameter. All transformations were performed separately in both bound states and in bulk bilayer to create complete thermodynamic cycles. In this study, each transformation was split into 15 windows where Coulombic and Van der Waals interactions were transformed separately along the λ co-ordinate, with soft-core parameters (α = 0.5 and σ = 0.3) for both interactions. Coulombic interactions’ λ parameter was perturbed linearly in the first 9 windows remains at one until the end (λ = 0.00, 0.10, 0.20, …, 1.00), while Van der Waals interactions’ λ parameter was perturbed linearly started from the sixth window toward the final window (λ = 0.00 … 0.00, 0.10, 0.20, …, 0.90, 1.00). Energy minimizations were performed with the steepest descent algorithm for 200 steps. 15 independent production simulations with randomized initial velocities were then run using leap-frog stochastic dynamics integrator to 12 ns per window, where the first 2 ns were discarded as equilibration. Using the alchemical-analysis software package, we constructed free energy pathways from individual simulation windows with Multistate Bennett Acceptance Ratio (MBAR)^86^.

### All-atom MD simulations

Structures were then converted from coarse-grained to all-atom representation using CG2AT2^87^. Specifically, H73 on KCNE1 was protonated to reflect its binding in the previous study^33^. The system was then energy minimized using steepest-descent algorithm for 5000 steps and then equilibrated for 10 ns with a restraint of 1000 kJ mol^-1^ nm^-2^ on the Cα atoms. All simulation temperatures were maintained at 310K using a v-rescale thermostat and at 1 bar semi-isotropic pressure coupling using a c-rescale barostat^88^. Three 500 ns production simulations for each conformation were conducted with different randomized initial velocities. All all-atom simulations were performed using GROMACS 2021.3 with the CHARMM36m forcefield.

### PIP2 titration using inside-out patch clamp electrophysiology

Inside-out patch clamp experiments were performed at room temperature using an Axon 200 B amplifier and an Axon^TM^ Digidata 1500 B digitizer. Pipette resistance was maintained between 0.5 and 1.5 MΩ. For KCNQ1–KCNE1 channels, the pipette solution comprised (in mM): 140 KOH, 5 EGTA, 2 KCl, and 20 HEPES, with pH adjusted to 7.2 using NaOH. The pipette solution for KCNQ1-KCNE3 channels included (in mM): 96 NaCl, 2 KCl, 1.8 CaCl2, 1 MgCl2, and 5 HEPES, with pH adjusted to 7.4 using NaOH. The internal solution contained (in mM): 140 KOH, 5 EGTA, 2 KCl, 20 HEPES, and 5 Mg-ATP, with pH set to 7.2 using NaOH. PIP2 (Echelon, P-4508) was dissolved in the internal solution, stored at -80°C, diluted to working concentration, and sonicated prior to use. For KCNQ1–KCNE1 channels, the protocol involved holding at -80 mV, a test potential of +40 mV, and a return to -80 mV. For KCNQ1–KCNE3 channels, the protocol used a holding potential of -60 mV, a +40-mV depolarization, and a return to -60 mV. Data analysis was performed using Clampfit 11.2.

### GPCR sensitivity and mutation analysis in oocytes

#### Channel expression in *Xenopus* oocytes

Stage V or VI oocytes were obtained from *Xenopus laevis* by laparotomy in accordance with the protocol approved by the Washington University Animal Institute Animal care and Use Committee Office. Oocytes were digested by collagenase (0.8 mg/mL, Sigma-Aldrich, MO) and injected with channel cRNAs (Drummond Nanoject, PA). Each oocyte was injected with 9.2 ng WT or mutant KCNQ1. For experiments with KCNE1 or KCNE3 expressions in Figures 1A, 2B, S1B, S7A-D and S10, KCNQ1 and KCNE1 or KCNE3 were co-injected at 4:1 (KCNQ1/KCNE1 mutants: KCNE1) and 2: 1 (KCNQ1/KCNQ1 mutants: KCNE3/KCNE3 mutants) mass ratio, respectively. For experiments with M1R expression in Figure 4B and S9, M1R was co-injected at 10:1 (KCNQ1–KCNE1/3: M1R). Injected oocytes were incubated in ND96 solution (96 mM NaCl, 2 mM KCl, 1.8 mM CaCl_2_, 1 mM MgCl_2_, 1 mM MgCl_2_, 5 mM HEPES) supplemented with 2.5 mM CH_3_COCO_2_Na, and 1:100 penicillin-streptomycin (pH 7.6) at 18 °C for 2-4 days before recording.

#### Two-electrode voltage clamp

Microelectrodes were pulled with borosilicated glass (B150-117-10, Sutter Instrument, CA) with a micropipette puller (P-1000, Sutter Instrument, Novato, CA) to a resistance of 0.3 – 3 MΩ when filled with 3M KCl solution and submerged in ND96 solution at room temperature. Whole-cell currents were recorded with a GeneClamp 500B amplifier (Axon instrument, CA) driven by Patchmaster (HEKA, Holliston, MA). To avoid aliasing, the device applied a low-pass filter to the measured currents at 2 kHz. In drug application experiments, involving *m*-3M3FBS (Sigma Aldrich, MO), Oxo-M (Tocris Bioscience, Bristol), and XE991(Sigma Aldrich, MO), appropriate amount of stock drug was directly added to the bath solution by a manual pipette to the desired final concentration. The recording chamber was thoroughly rinsed with 70% ethanol the deionized water after each experiment involving drug application.

#### Data analysis

Conductance-voltage (*GV*) relationship was conducted by measuring the instantaneous tail currents following test pulses, which were normalized to the maximum tail current and fitted with the Boltzmann function in the form of:

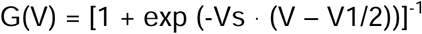

where V is the test voltage, Vs is the slope, and V_1/2_ is the voltage at half of maximum Gmax.

## Notes

### Competing Interest Statement

The authors have declared no competing interest.

### Summary of Updates

1. References 1 and 2 are removed from Abstract. 2. Affiliation of Washington University in St. Louis is uniformed. 3. Affiliation of KTH is detailed as KTH Royal Institute of Technology

