## Supplementary figures and images for "Mechanisms of KCNQ1 gating modulation by KCNE1/3 for cell-specific function"

### Supplemental Figure 1, and will be used for the link to the file on the preprint site

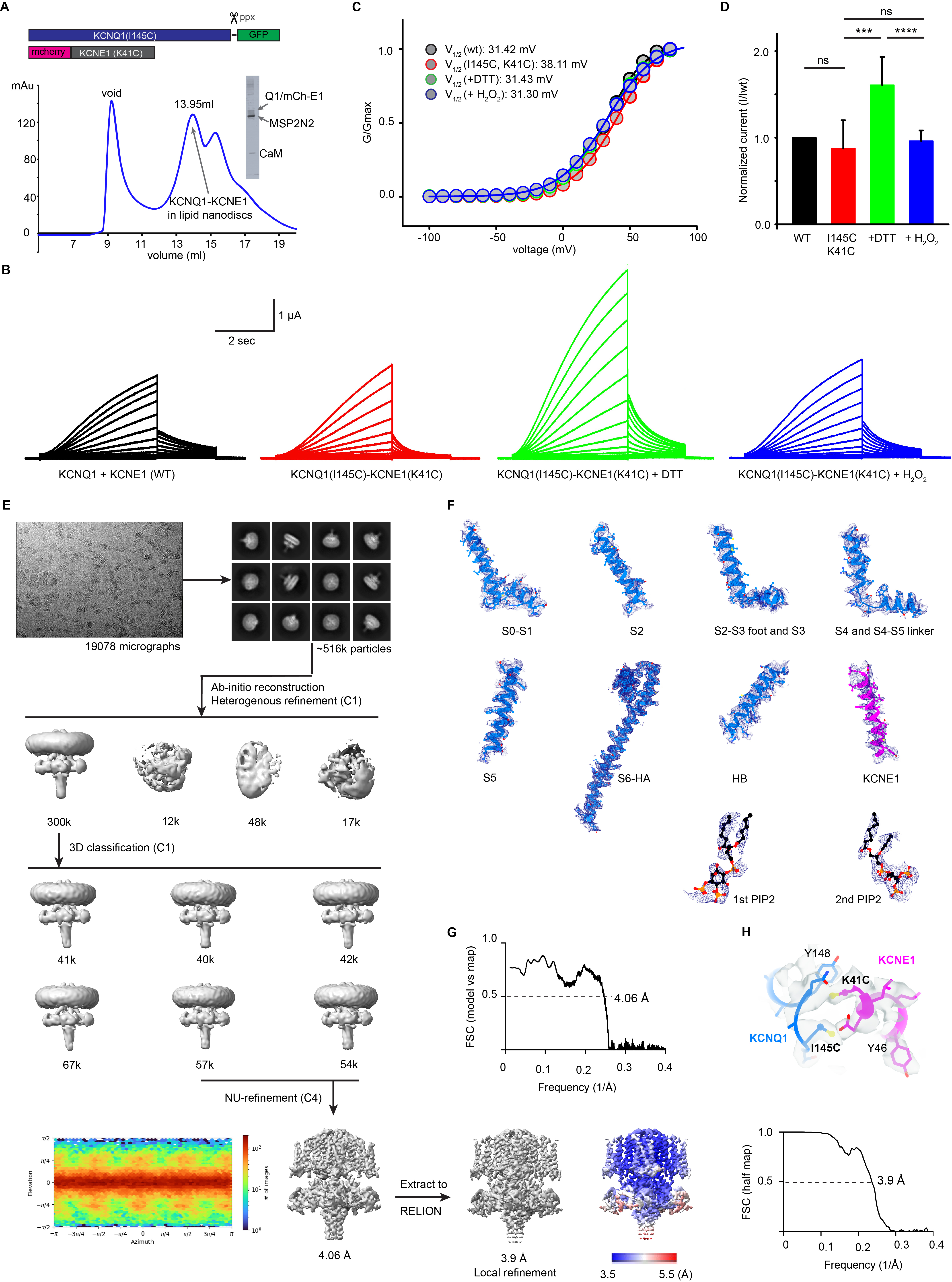

### Supplemental Figure 2, and will be used for the link to the file on the preprint site

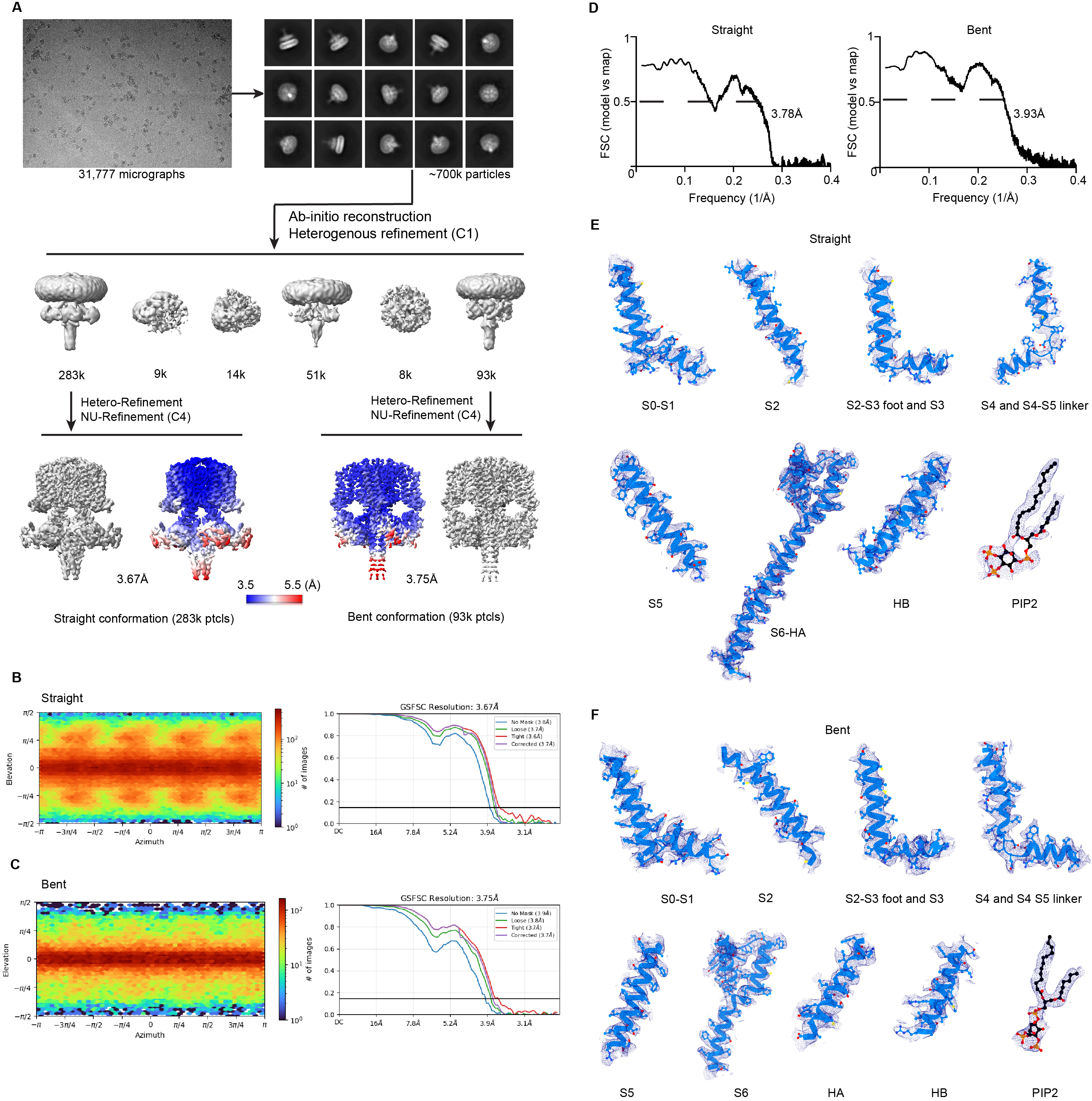

### Supplemental Figure 3, and will be used for the link to the file on the preprint site

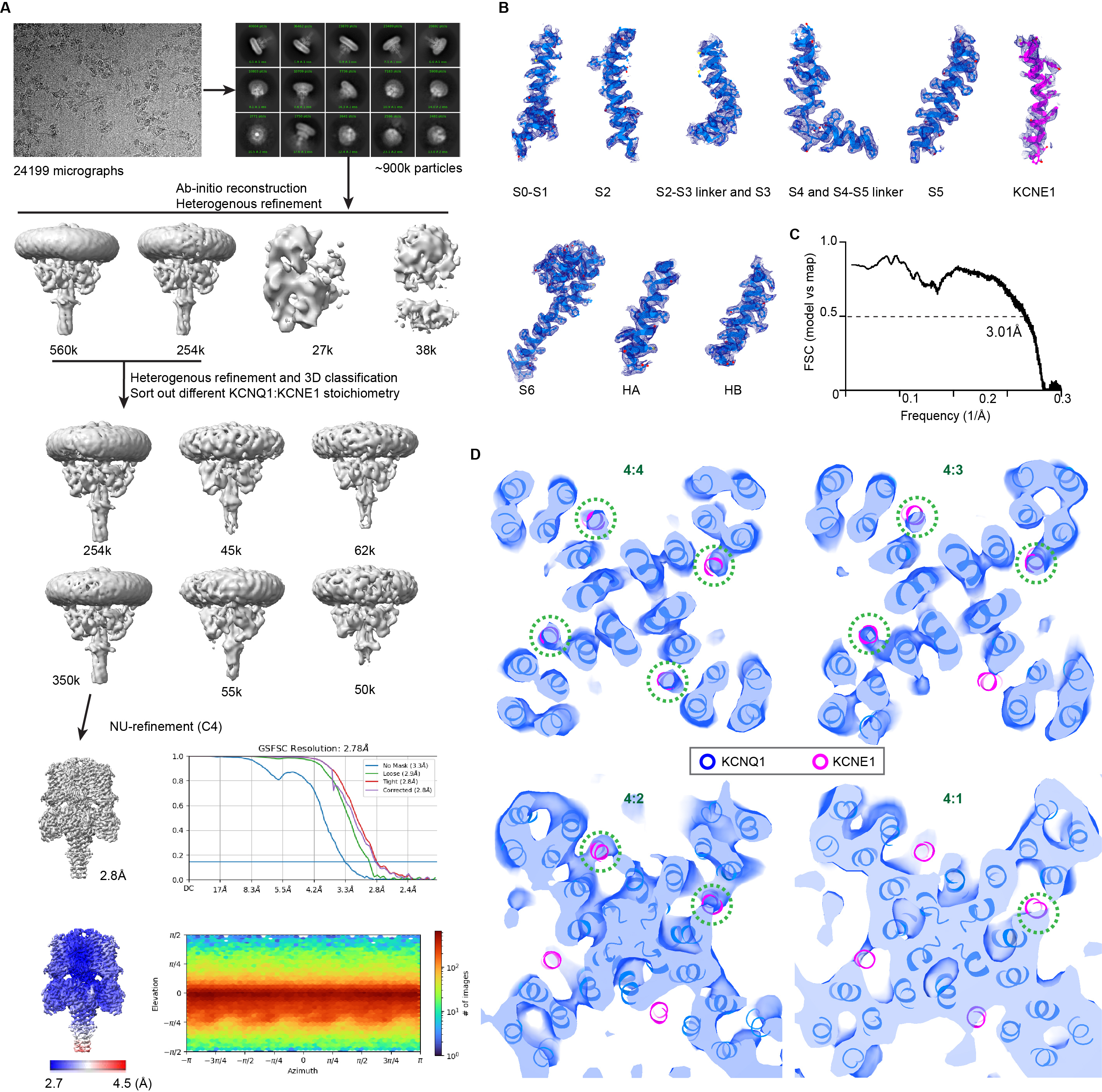

### Supplemental Figure 4, and will be used for the link to the file on the preprint site

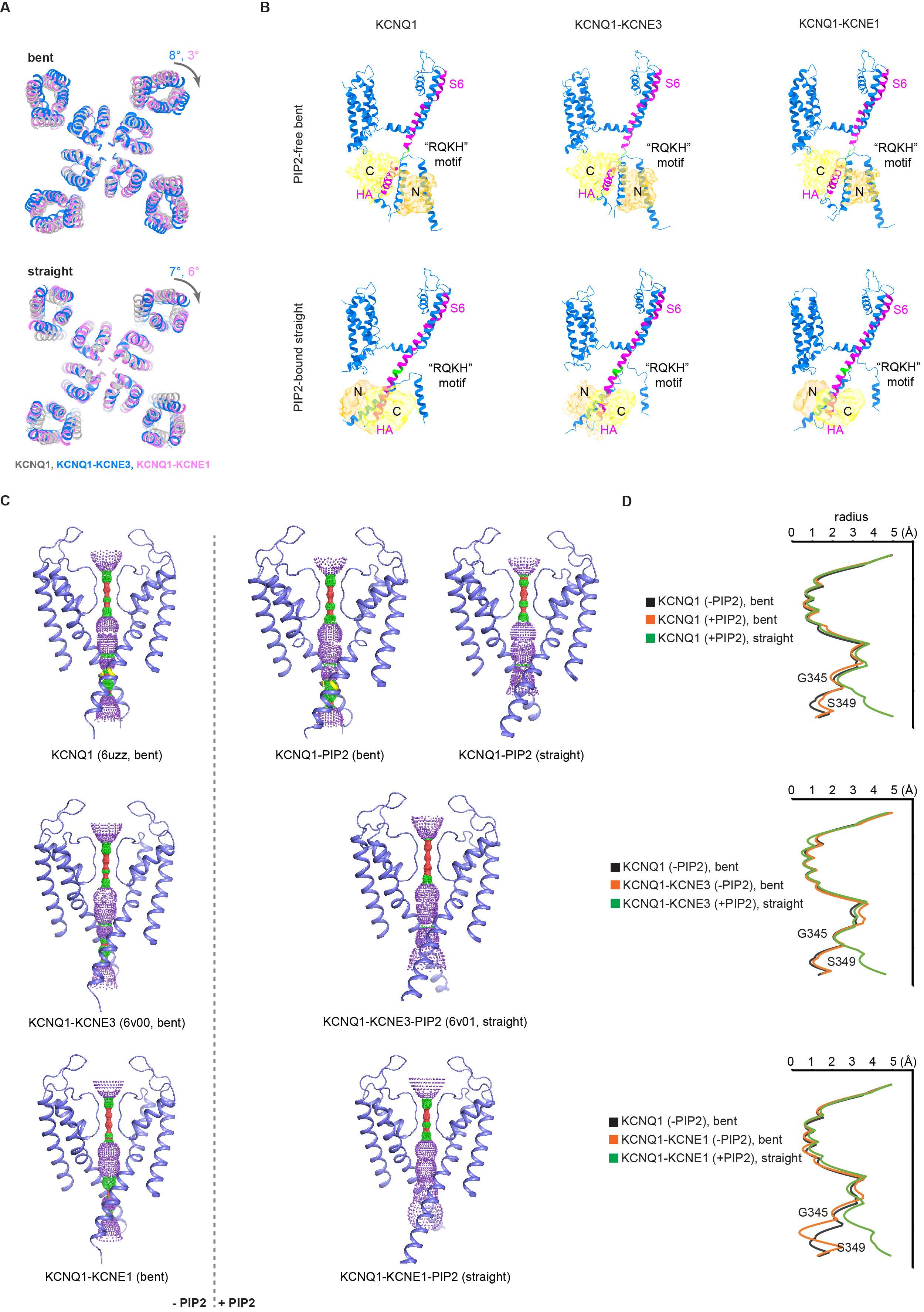

### Supplemental Figure 5, and will be used for the link to the file on the preprint site

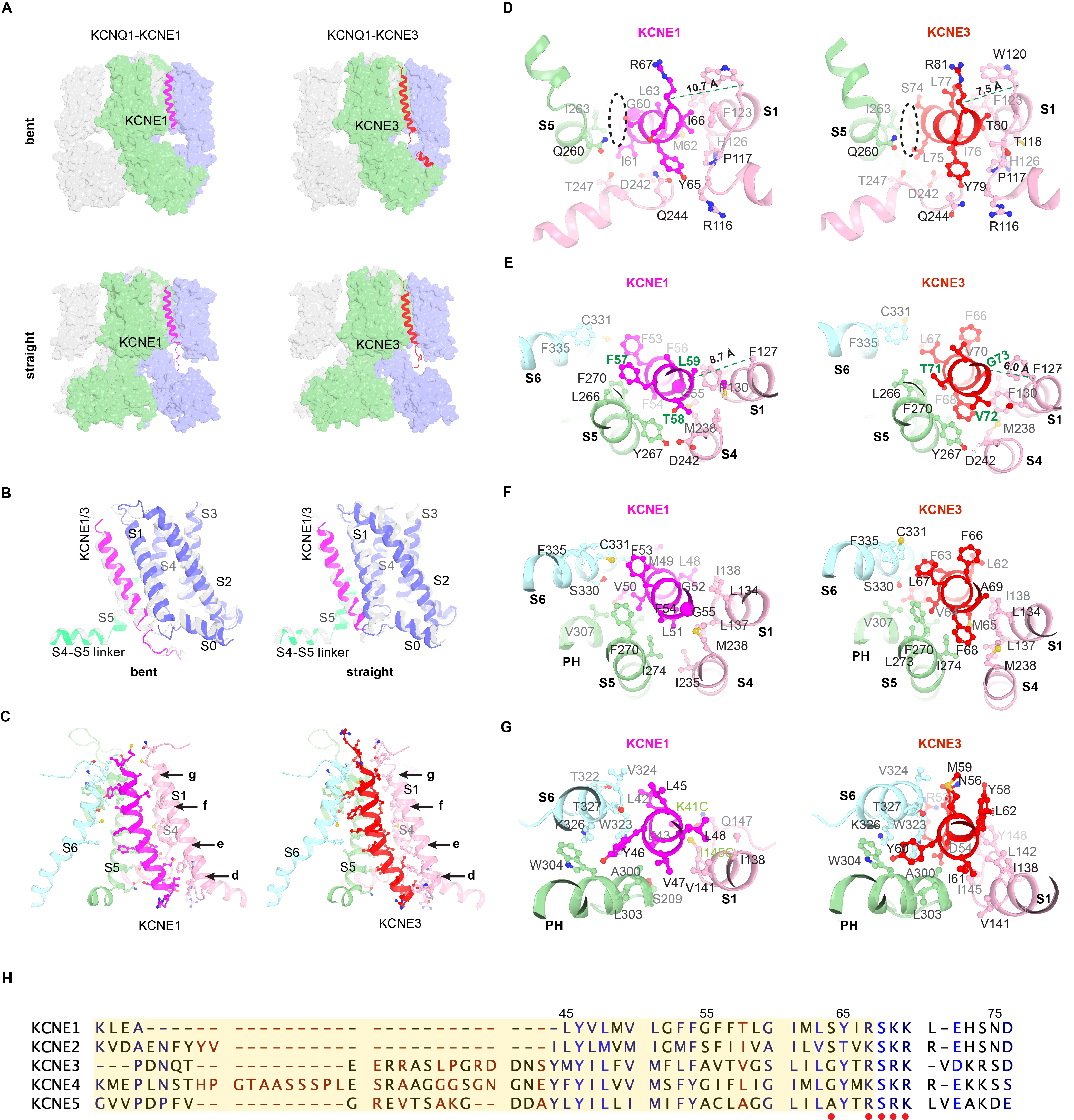

### Supplemental Figure 6, and will be used for the link to the file on the preprint site

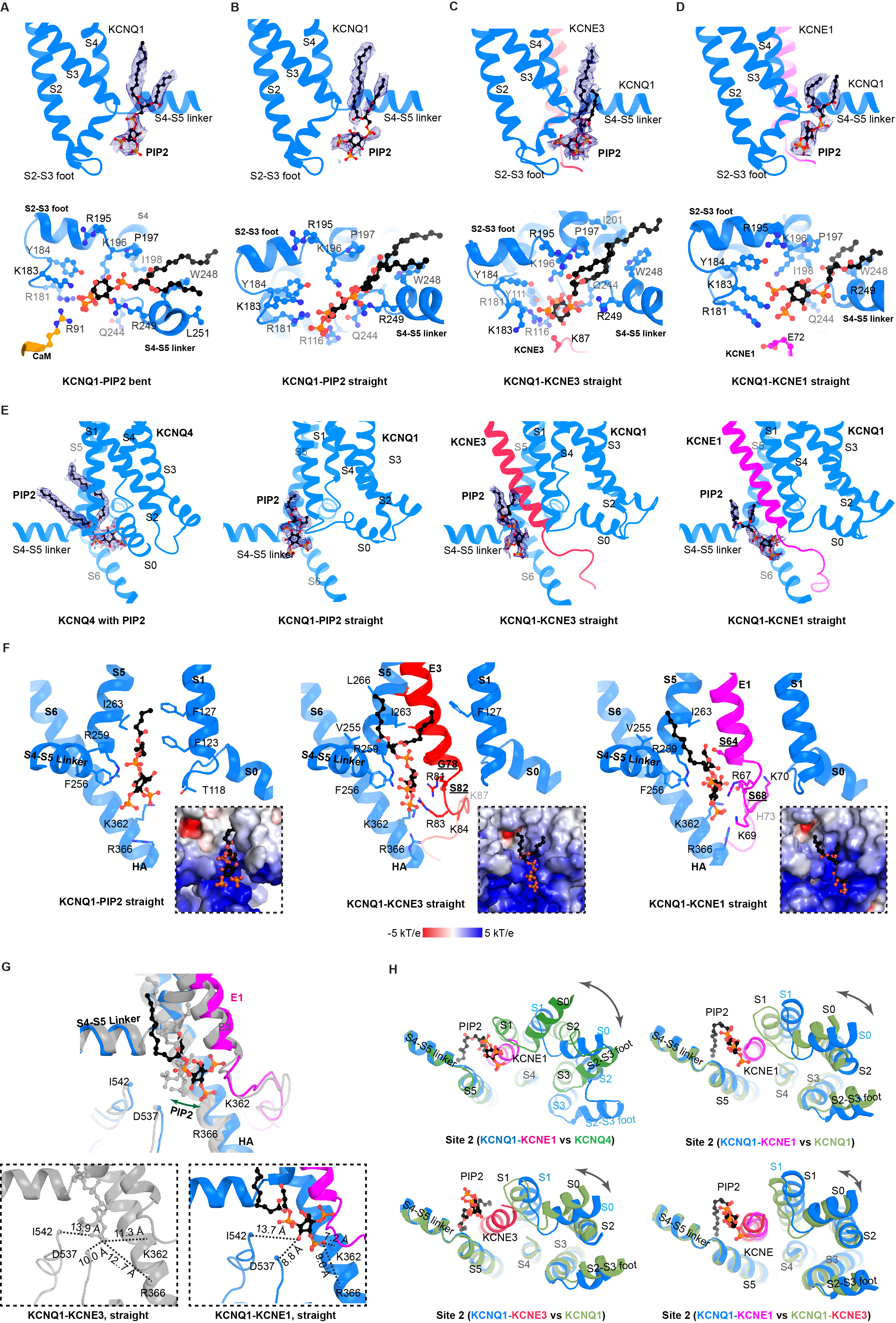

### Supplemental Figure 7, and will be used for the link to the file on the preprint site

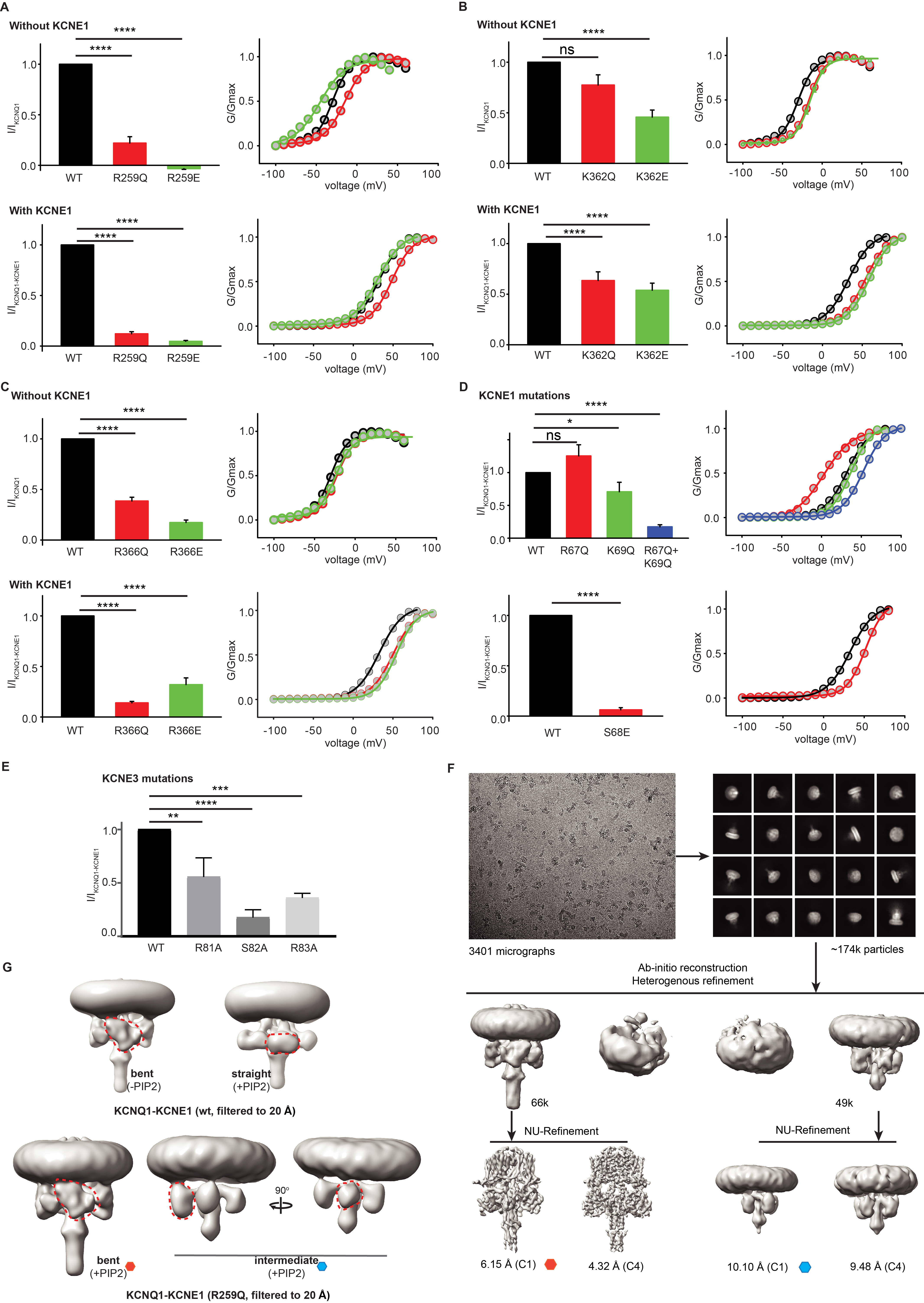

### Supplemental Figure 8, and will be used for the link to the file on the preprint site

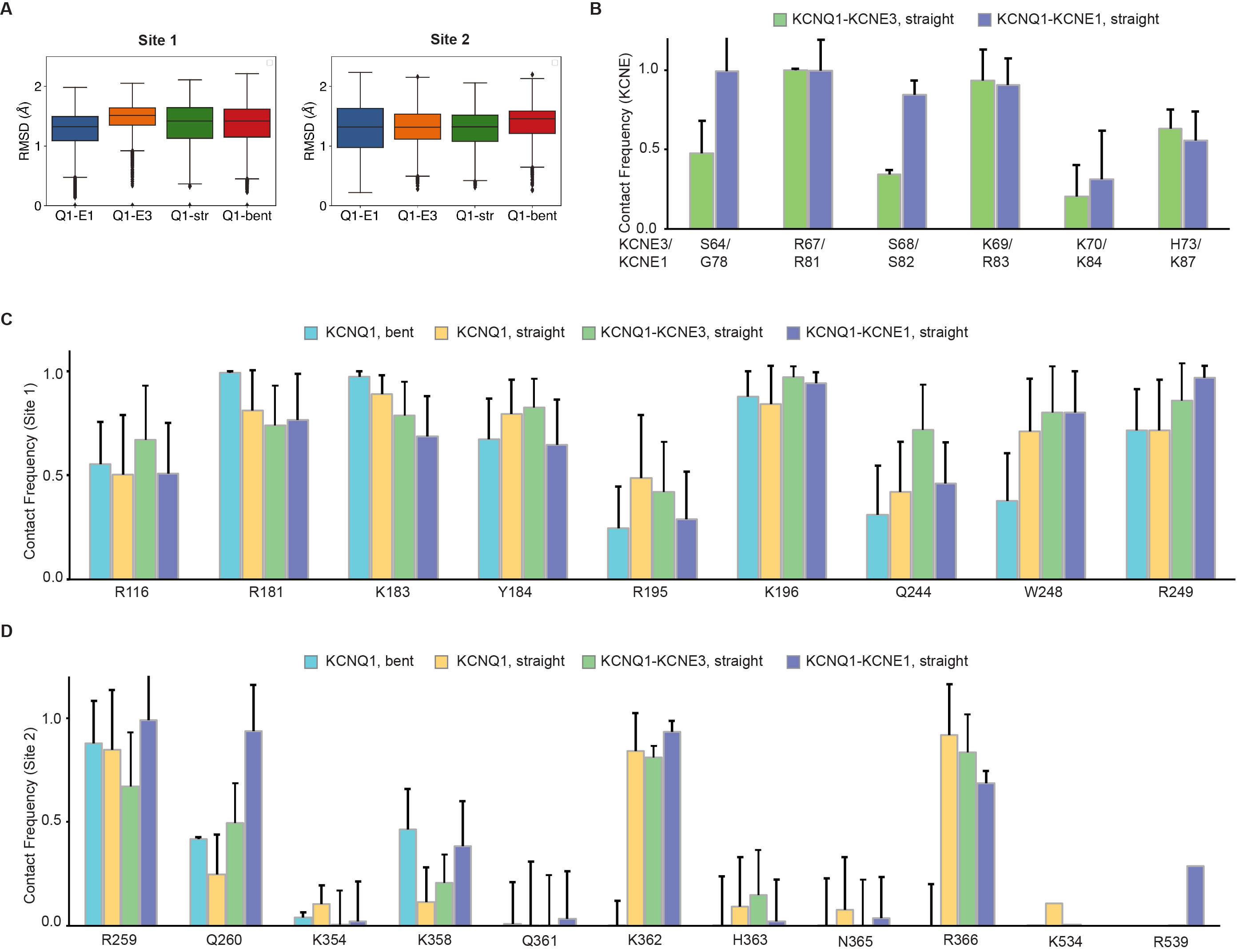

### Supplemental Figure 9, and will be used for the link to the file on the preprint site

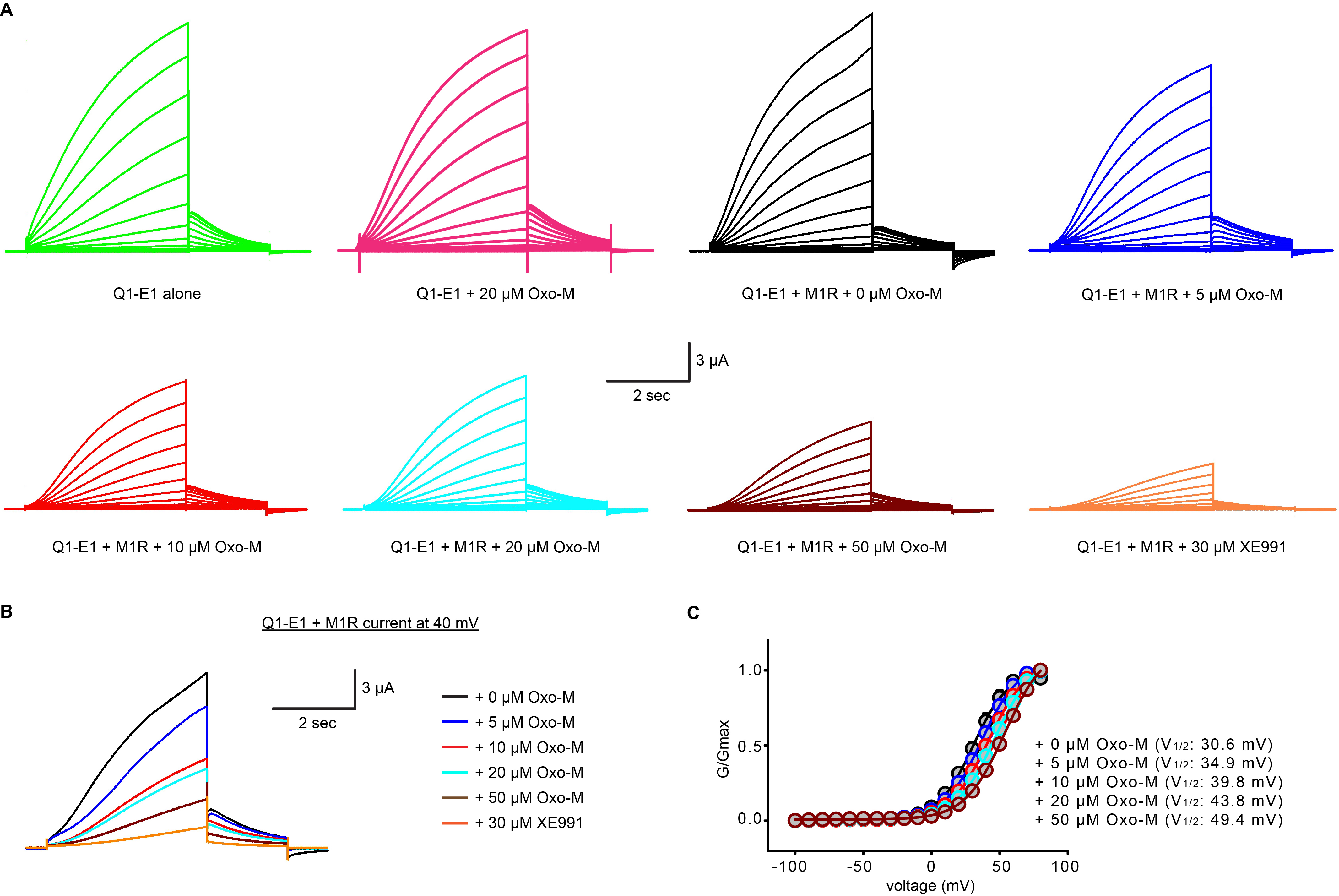

### Supplemental Figure 10, and will be used for the link to the file on the preprint site

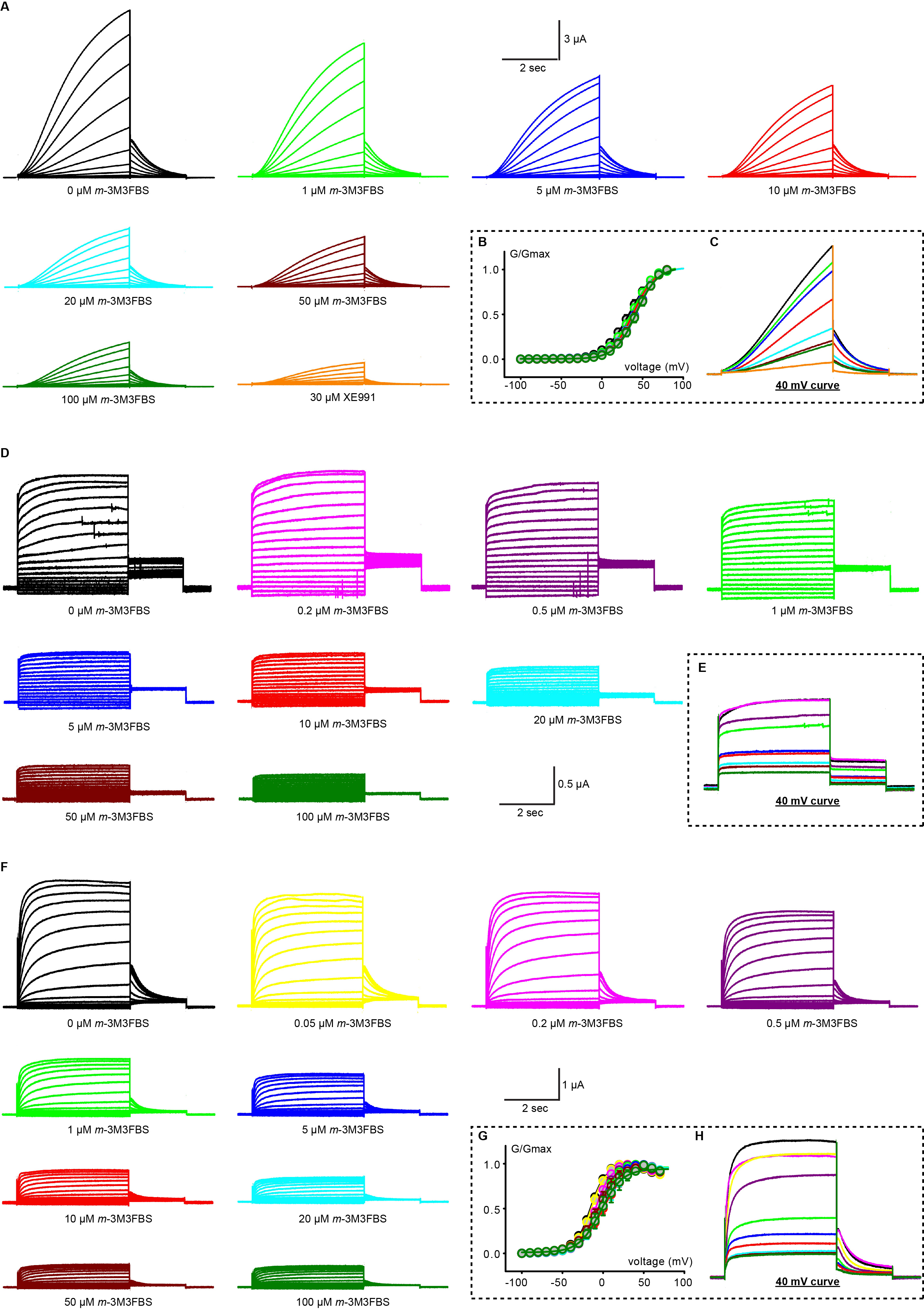
